# Multi-omics analysis identifies intrinsic *Trp53* driven metastatic breast cancer subtypes

**DOI:** 10.64898/2026.02.16.706177

**Authors:** Joy M. McDaniel, Rhiannon L. Morrissey, Gilda P. Chau, Xiaoping Su, Zach Von Ruff, Anil Korkut, Vidhi Chandra, Lei Huo, Adel K. El-Naggar, Guillermina Lozano

**Author notes:** **Corresponding Author**: Guillermina Lozano.

## Abstract

Metastatic breast cancer (mBC) is deadly, and its molecular drivers are largely unknown. Treatment is limited to systemic cytotoxic chemotherapies and tumors often become refractory. The most frequently mutated gene in MBC is *TP53*. The *TP53^R248W^* hotspot missense mutation is associated with poor prognosis. We generated a somatic mouse model of mammary epithelial specific *Trp53^R245W^*expression. Primary tumors were highly metastatic, reflecting the human molecular subtypes luminal A, luminal B, HER2, and TNBC. Transcriptomic profiling revealed three intrinsic subtypes: stem-cell like (SCL), well-differentiated metabolically active (WDMA), and immunosuppressed (IS). SCL tumors activate ribosome biosynthesis and E2F signaling, amplifying *Met*, *Birc3*, *Yap1* and deleting *Nf1*, *Pik3r1*, and *Rad17*. WDMA tumors activate cytochrome P450 enzymes, estrogen signaling and branched chain amino acid degradation, with mutations activating Pi3k/Akt/mTOR signaling. IS tumors activate immune suppression, have high mutation burden, and frequently mutate *Traf7* and delete *Cdkn2a*. This is the most comprehensive transcriptomic and genomic profiling of mutant p53-driven breast tumors, elucidating potential therapeutic targets.

**Teaser:** A single *Trp53* mutation drives intrinsic metastatic breast cancer subtypes with distinct transcriptomes and genomic alterations.

## Introduction

Breast cancer is the most frequently diagnosed cancer in women worldwide. Seventy to eighty percent of patients diagnosed with early-stage, non-metastatic disease, respond to therapy and are cured (*1*). However, 30% of all breast cancer diagnoses become advanced with distant organ metastases, referred to as metastatic breast cancer (mBC) and is incurable (*2*). Ninety percent of all cancer deaths are due to metastatic disease (*3, 4*). Current treatment for mBC is cytotoxic chemotherapy combined with hormone therapy, targeted anti-HER2 therapy, or immunotherapy depending on the hormone receptor (HR) status of the tumor. Two-year survival rates vary depending on the molecular subtype: (1) 45.5% HR positive mBC, HER2 positive and/or negative (luminal B) mBC (2) 35.9% HR positive, HER2 negative (luminal A) mBC; (3) 33.9% HR negative, HER2+ (HER2-enriched) mBC; and 11.2% triple negative breast cancer (TNBC) (*5*). mBCs are often treatment-resistant, resulting in death. It is critical to understand the molecular underpinnings of mBC to inform development of targeted therapies for this disease.

Next generation sequencing has revealed the most frequently altered gene in mBC is *TP53* (*6, 7*)*. TP53* alterations are associated with endocrine resistance in luminal breast cancers, targeted therapy resistance in HER2-enriched breast tumors; and chemoresistance in TNBCs (*8–14*). *TP53* mutations occur in 12% luminal A; 29% luminal B, 72% HER2-enriched; and with the highest frequency in 88% TNBC (*15*). *TP53* mutations are early driver events contributing to initiation and progression.

The *TP53* tumor suppressor gene encodes a transcription factor that is present at low levels until activated in response to cellular and genotoxic stressors including DNA damage, oxidative stress, and oncogenic mutations (*16, 17*). Tumor suppressors are often deleted in many cancers. Contrary to other tumor suppressors, most *TP53* alterations in cancer are missense mutations, present at hotspot amino acids of the DNA-binding domain. *TP53* hotspot mutations frequently undergo loss of heterozygosity and often impart various oncogenic advantages to the cell, including enhanced metastatic potential (*18–21*). In breast cancer, the hotspot *TP53^R248W^* missense mutation is associated with poor prognosis, and mBC.

To delineate the *in vivo* role of mutant p53 in the development and progression of mBC, we have developed a physiologically relevant somatic mouse model, in which the *Trp53^R245W^*missense mutation (orthologue to the human breast cancer hotspot missense mutation *TP53^R248W^*), is induced temporally and spatially. Conversion of wild-type (WT) to mutant p53 in the mammary gland via intraductal delivery of adenovirus containing Cre-recombinase (referred to as Ad-Cre) results in excision of WT p53 cDNA sequences from the endogenous locus, resulting in <u>ma</u>mmary ductal epithelial cell-specific mutant <u>p</u>53 expression of *Trp53^R245W^*(*MaP^R245W^* from hereon), with preservation of WT p53 in the stroma and immune system (*22–24*). The ability to mimic a single somatic p53 mutation in the murine mammary duct, with wild type immune and stroma, with limited genetic variation, allows us to understand how mutant p53 influences the genome, transcriptome and immune microenvironment to mediate mBC onset and progression. *MaP^R245W^* mice develop mBC. Primary tumors from this *MaP^R245W^* model reflect the human molecular breast cancer subtypes, including luminal A, luminal B, HER2-enriched, and TNBC (*22–24*). Most tumors were of the TNBC subtype, making our model an excellent source for modeling TNBC, since 88% of human tumors of this subtype harbor a *p53* alteration.

Comprehensive integrated omics data are urgently needed to shed light on the role of an initiating driver mutant p53 in mBC, particularly the dysregulated pathways and cooperating genomic events associated with it. Here we present a large cohort of mice from our *MaP^R245W/+^* driven somatic model of mBC, for which we have comprehensive multi-omics analyses via bulk RNA-sequencing and whole exome sequencing of primary mammary tumors. Integration of transcriptomic and genomic data revealed three distinct molecular subtypes amongst *MaP^R245W/+^*mammary tumors with varied intrinsic oncogenic factors. We investigated human breast cancers harboring *p53* mutations, and identified similar intrinsic signatures, corroborating our model mimics human mBC. Our study provides insight into the biology of metastatic breast cancers harboring a p53 mutation, underscores the importance of considering p53 mutation status in breast cancer and provides a resource for future development of precision medicine approaches for mBC.

## Results

### A Somatic Model of Metastatic Breast Cancer

The *TP53^R248W^* mutation (human orthologue of mouse *Trp53^R245W^*) is the most frequent hotspot mutation in breast cancer and correlates with poor prognosis and reduced survival compared to other *TP53* mutations (*25*). To model how the R245W missense mutation facilitates mammary tumor development and progression, we generated *MaP^R245W/+^* mice (n=80 mice) via intraductal delivery of adenovirus containing Cre recombinase (Ad-Cre) to the epithelial cells of the mammary duct. Intraductal delivery of Ad-Cre resulted in recombination at one WT *Trp53* allele to express mutant p53. In this model, p53R245W was expressed specifically in epithelial cells, while the stroma and the immune environment retained WT *Trp53* (*22–24*). Analysis of *MaP^R245W/+^* mice showed 61% developed primary mammary tumors (49 of 80 mice), at endpoint, with a mean survival of 22.5 months (Fig. 1A, Supplementary Table 1). Approximately, 43% (34 of 80 mice) of mice in this cohort harbor the Tdtomato reporter allele, allowing us to trace the recombination and metastatic dissemination of tumor cells (Table S1). In a separate cohort, we have previously demonstrated the presence of the Tdtomato reporter does not result in differences in survival as compared to mice that do not harbor the reporter (*24*). Additionally, we have performed time course studies in *MaP^R245W/+^*; *Rosa26^LSL-TdTomato^* mice at several time points post Ad-Cre injection to test Cre leakiness. IVIS imaging of mice at 1, 3-, 7-, 10-, and 12-months post-injection revealed *TdTomato* fluoresces only in the mammary fat pads, suggesting recombination occurs only in the mammary duct (Fig. S1). These data indicate the presence of a *Trp53* mutation in the mammary duct is sufficient to initiate tumorigenesis and this model is useful to identify transcriptomic and genomic changes associated with this mutation.

**Fig. 1.**
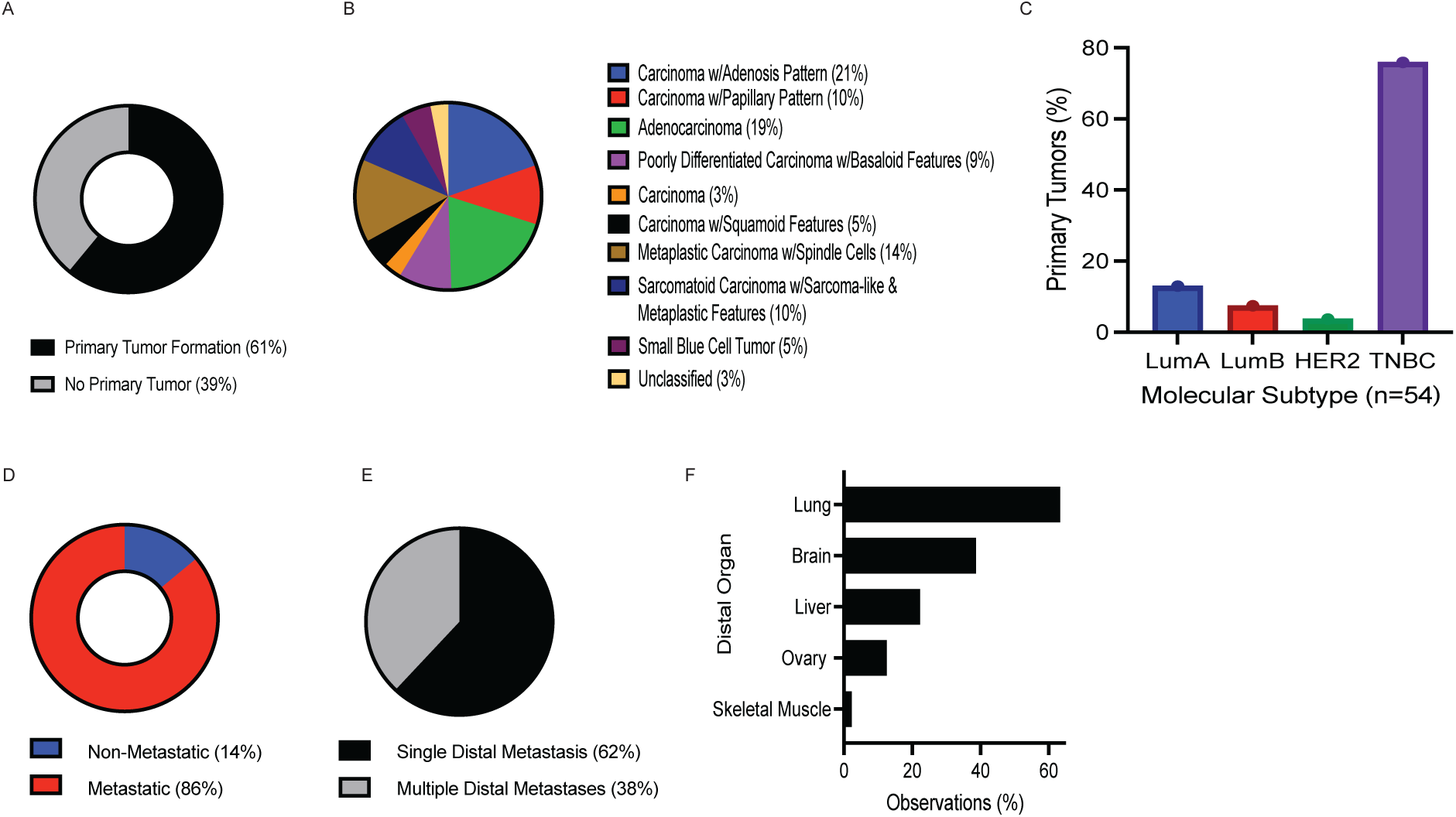
Molecular Characterization of Mammary Tumors from *MaP^R245W/+^* Mice. (A)Incidence of primary tumor formation and mice without primary tumor formation in *MaP^R245W/+^* cohort (B) Pie chart depicting the histological subtypes of the mammary tumors from *MaP^R245W/+^*mice. (C) Breast cancer molecular subtypes defined by qPCR for *Esr1* (Estrogen Receptor), *Pgr* (Progesterone Receptor), and *Erbb2* (HER2). (D) Incidence of non-metastatic and metastatic mice. (E) Incidence of single and multiple metastatic events in mice with primary mammary tumors. (F) Organs with metastatic lesions observed.

Histological analysis was performed on primary mammary tumors from *MaP^R245W/+^* mice (n=49). Six of these mice developed two mammary tumors and two tumors had a bi-lobed appearance for which individual cuts were taken. Thus, fifty-eight mammary lesions were assessed, which includes multiple tumors from individual mice and bi-lobed tumors. Except for five lesions, all mammary lesions were carcinomas (Table S1). The carcinomas comprised eight histological categories: (1) carcinomas with adenosis pattern; (2) adenocarcinoma; (3) metaplastic carcinoma with spindle cells; (4) carcinomas with papillary pattern; (5) poorly differentiated carcinoma with basaloid features; (6) sarcomatoid carcinoma with sarcoma-like and metaplastic features; (7) carcinoma with squamoid features; and (8) carcinoma, not classified (Fig. 1B, Fig. S2A, Table S1). Interestingly, carcinomas with adenosis or papillary patterns, represent a class of well-differentiated carcinomas that have retained their ductal structures, characterized by tight packing of glands (adenosis) or small duct protrusions (papillary). Tumors classified as metaplastic with spindle cell, poorly differentiated with basaloid and sarcomatoid carcinoma with sarcoma-like and metaplastic features, each represent rare high-grade human breast tumors (*26*).

Three mammary lesions were classified as small blue cell tumors (Fig. 1B, Fig. S2A, and Table S1). Small blue cell tumors represent a group of undifferentiated aggressive embryonal tumors that include small desmoplastic round blue cell tumors; which are rare in humans but do occur in the breast and are often metastatic (*27*) (*28*). Two tumors were de-differentiated to the point they were denoted as unclassified (Fig. 1B, Fig. S2A, and Table S1). Collectively, the preferential development of aggressive breast tumors in this *MaP^R245W/+^* model reiterates findings demonstrating human breast cancers with *TP53* mutations reflect an aggressive disease state and that the human *TP53^R248^* missense mutation is associated with poor prognosis in breast cancer (*25, 29, 30*).

We have previously demonstrated that our somatic model mimics human breast cancer molecular subtypes (*22–24*). Real time qRT-PCR analysis was conducted to assess the expression levels of *Esr1* (estrogen receptor), *Pgr* (progesterone receptor), and *Erbb2* (human epidermal growth factor receptor2, HER2), in all primary mammary tumors from *MaP^R245W/+^* mice, except for four lesions (Table S1). These four lesions had undetectable transcript levels by qRT-PCR experiments and were thus labeled as unclassified for their molecular subtype. A total of 54 primary mammary lesions were analyzed. Most *MaP^R245W/+^* mammary tumors are of the TNBC subtype, with 76% lacking expression of *Esr1*, *Pgr*, and *Erbb2* expression (Fig. 1C, Table S1). Luminal A (*Esr1^+^*, *Pgr^+^*) disease represented 13% of the mammary lesions (Fig. 1C, Table S1). Luminal B lesions are characterized by positive expression of *Esr1* and/or *Pgr* and are either *Erbb2* positive or negative. The luminal B subtype comprised 7% of *MaP^R245W/+^* mammary lesions (Table S1, Fig. 1C). Lastly, 4% of mammary tumors mimicked the HER2-enriched subtype, characterized by only *Erbb2* expression (Fig. 1C, Table S1). These results demonstrate that our *MaP^R245W/+^* somatic model of mBC develops primary tumors reflecting the human molecular subtypes of breast cancer.

### MaP^R245W/+^ Mice are Highly Metastatic

*Most MaP^R245W/+^* mice develop a metastatic phenotype. Among the 49 mice with primary tumor formation, 86% (42 of 49 mice) developed distal metastases (Fig. 1D, Table S1). Metastases were identified by gross dissection, histopathology, and visualization of the TdTomato fluorescent reporter, when present (Table S1). 62% of metastatic mice had a single organ metastasis (26 of 42 mice), while 38% (16 of 42 mice) had multi-organ metastases (Fig. 1E, Table S1). Distant metastases were observed in the following organs: lung (63%), liver (22%), brain (8%), ovary (5%), and skeletal muscle (2%) (Fig. 1F, Table S1). These findings suggest our model closely mimics human metastatic breast cancer, as it preferentially metastasizes to organs frequently observed in humans. Additionally, the high prevalence of TNBC in this model corroborates the observation that 88% of TNBCs harbor a *TP53* alteration and this tumor type is often metastatic. Also, the presence of luminal A, luminal B, and HER2-enriched subtypes in our model demonstrates the presence of a *p53* mutation induces a metastatic breast cancer phenotype regardless of the molecular subtype.

### A Single Trp53^R245W^ Mutation Induces Three Distinct Groups of Metastatic Breast Cancer

Large-scale genomics studies have identified *TP53* missense mutations in aggressive breast cancers including those that exhibit chemoresistance and metastasis (*31, 32*). Clonal evolution studies suggest *TP53* mutations likely initiate breast cancer, although there is no direct evidence (*33, 34*). *Trp53^R245W^ is* the initiating mutation in our somatic model of mBC, providing an opportunity to unveil intrinsic oncogenic signatures associated with mutant p53. To this end, we performed bulk RNA-sequencing on a total of 51 primary mammary lesions from *MaP^R245W/+^* mice (Table 1). Dimension reduction via principal components analysis (PCA) of RNA-sequencing data from *MaP^R245W/+^* mammary tumors revealed transcriptional heterogeneity across tumors (Fig. 2A). PCA analysis combined with k-means hierarchical clustering revealed 3 transcriptional subgroups: Groups A (n=28), B (n=13), and C (n=10) (Figs. 2A and B). *MaP^R245W/+^* tumor groups were not defined by metastatic phenotypes, histological subtypes, nor molecular subtypes (Figs. 2C and H-J, Figs. S3A-C). Principal components analysis revealed all tumors from the 8 non-metastatic mice cluster with Group A along with other tumors that are metastatic (Fig. 2C).

**Fig. 2.**
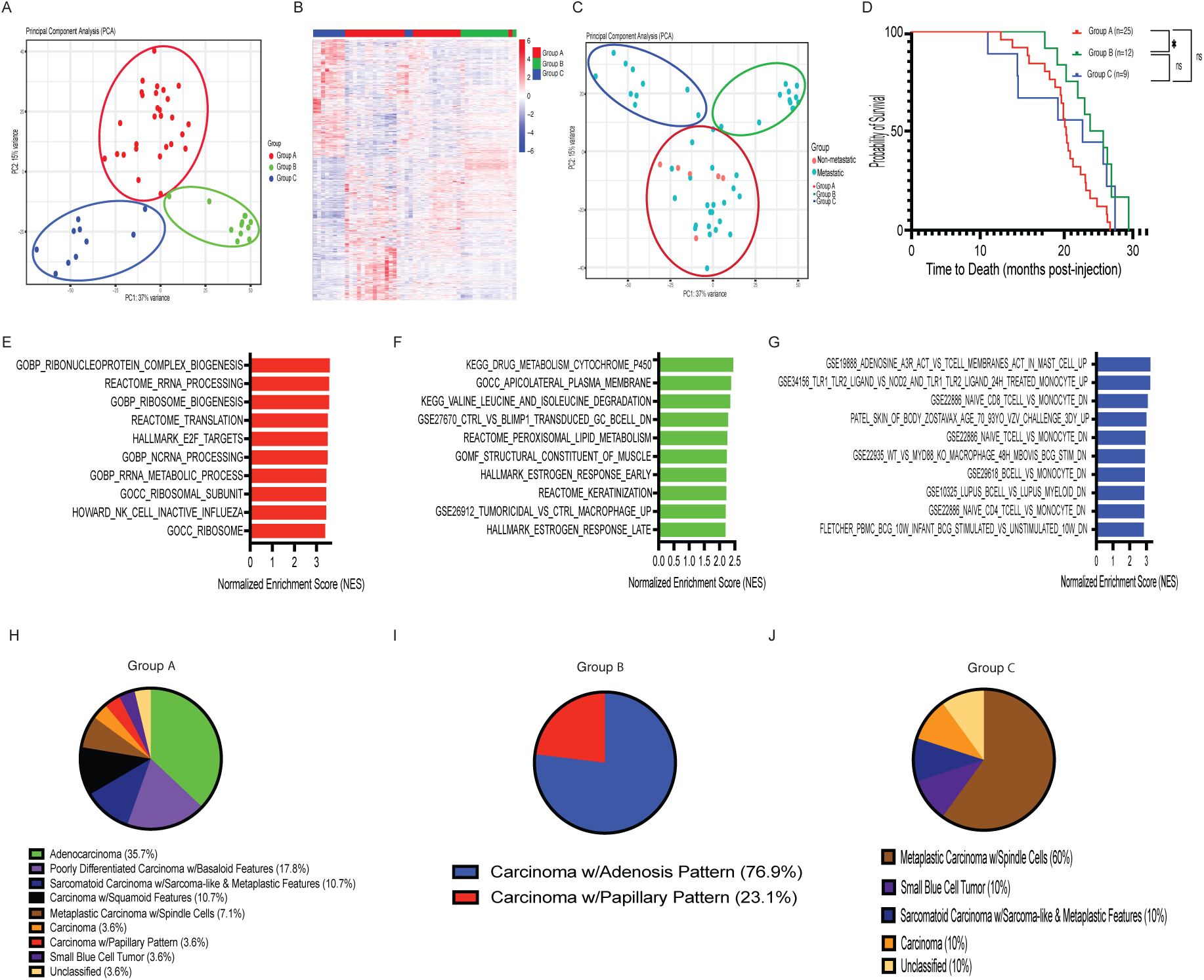
*MaP^R245W/+^* Mammary Tumors are Heterogeneous. (A) Principal components analysis depicting three transcriptional subgroups of *MaP^R245W/+^* tumors. (B) Heatmap showing unsupervised hierarchical clustering of *MaP^R245W/+^* and mammary tumors using the Pearson distance and Ward linkage. (C) Principal components analysis of transcriptomes from non-metastatic (pink) and metastatic (teal) mice from *MaP^R245W/+^* cohort. Colored circles indicate transcriptomic subgroup membership (Red=group A; Green=Group B; Blue=Group C). (D) Overall survival of Groups A, B, and C *MaP^R245W/+^* mice. (Intergroup survival comparison: Groups A vs B, p< 0.02; A vs C, ns; B vs C, ns) (E) GSEA results depicting significantly dysregulated pathways and processes in *MaP^R245W/+^* Group A mammary tumors. (F) GSEA results depicting significantly dysregulated pathways and processes in *MaP^R245W/+^* Group B mammary tumors. (G) GSEA results depicting significantly dysregulated pathways and processes in *MaP^R245W/+^* Group C mammary tumors. (H-I) Pie charts depicting the histological subtypes of mammary tumors from *MaP^R245W/+^* Groups A, B, and C mice, respectively.

**Table 1.**
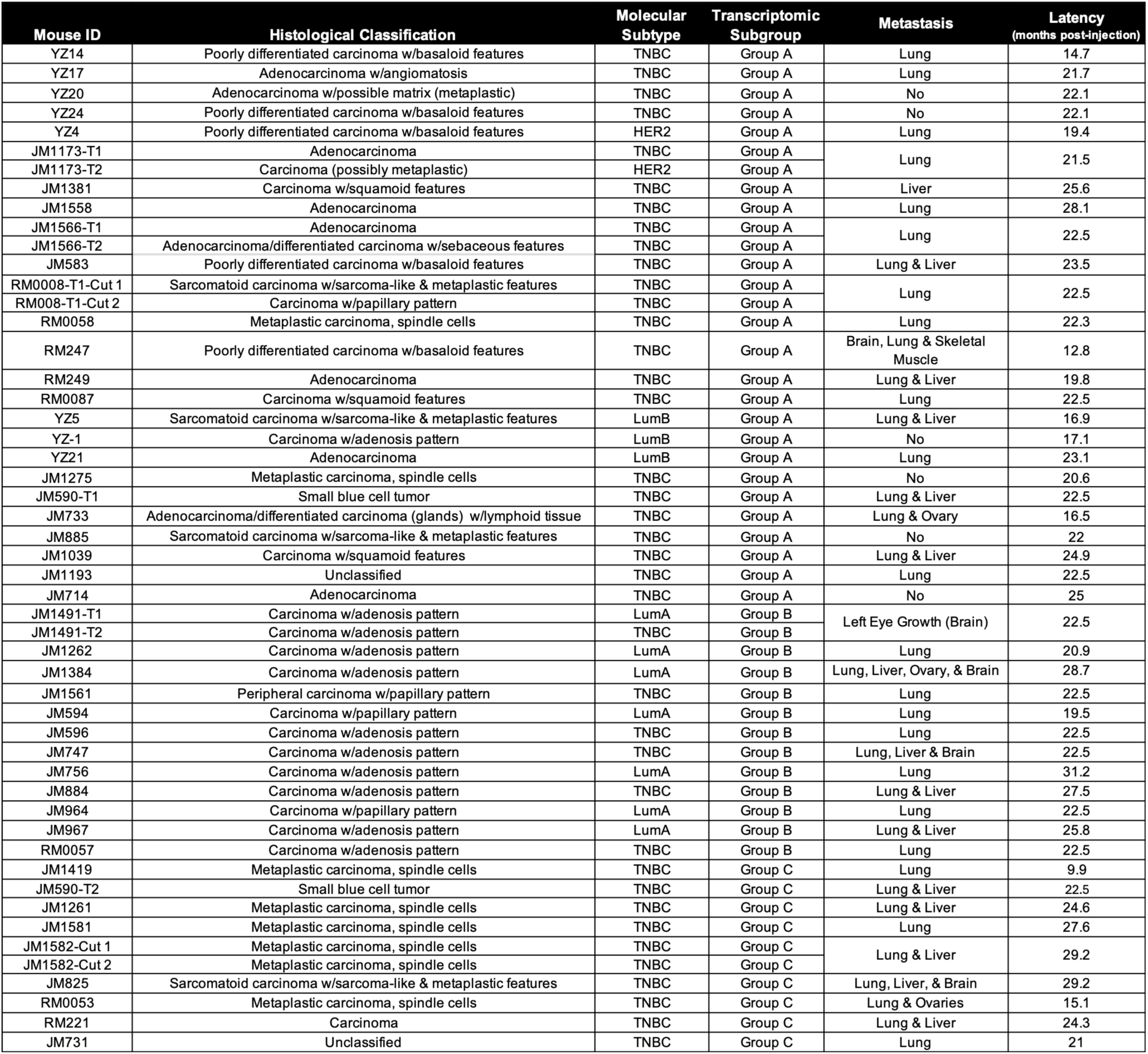
Transcriptomic Subgroup Assignments of *MaP^R245W/+^* Mammary Tumors. Summary of transcriptomic group membership for *MaP^R245W/^*^+^ mammary tumors based on principal components analysis.

Survival analysis was performed on mice from each transcriptomic group. Mice from Groups A, B, and C had a median latency of 22 months, 27 months, and 24 months, respectively. Intergroup survival analysis revealed statistically significant differences in survival between Group A mice compared to Group B mice only (p < 0.005) (Fig. 2D, Figs. S3D-F).

### Group A MaP^R245W/+^ Mammary Tumors Activate a Stem Cell Like Program

To examine the group specific transcriptomes contributing to the heterogeneity observed across *MaP^R245W/+^* mammary tumors, intergroup differential gene expression analysis was performed. Group A stem cell like tumors comprise a diverse group of rare aggressive histological classifications (n=28 tumors): adenocarcinomas, poorly differentiated carcinomas, sarcomatoid carcinoma with sarcoma-like and metaplastic features, carcinoma with squamoid features, metaplastic carcinoma with spindle cells, carcinoma, carcinoma with papillary pattern, and small blue cell tumor (Fig. 2H, Table 1). The following molecular subtypes identified in Group A mammary tumors (n=28) included: 82.1% TNBC, 7.1% HER2-enriched, and 10.7% Luminal B (Fig. S3A, Table 1).

Comparison of Group A mammary tumors (n=28) to Groups B and C tumors (n=23) revealed 3,089 differentially expressed genes (DEGs), of which 1,439 genes are upregulated, and 1,650 genes are downregulated using a false discovery rate (FDR) of 5% and log2 fold change of |1| (Figs. S3G and H). Gene Set Enrichment Analysis (GSEA) revealed significant enrichment in the following pathways and processes: GOBP Ribonucleoprotein Complex Biogenesis, Reactome RRNA Processing, GOBP Ribosome Biogenesis, Reactome Translation, Hallmark E2F Targets, GOBP NCRNA Processing, GOCC Ribosomal Subunit, Howard NK Cell Inactivation, and GOCC Ribosome (Fig. 2E) (*35–37*). Dysregulation of E2F signaling, ribosomal processes, and non-coding RNA processes not only contribute to epithelial to mesenchymal transition (EMT) and metastasis but are also phenotypic attributes of cancer stem cells. We will refer to Group A tumors as Stem Cell Like Tumors (*38–43*). Interestingly, TNBCs exhibit cancer stem cell like phenotypes more than non-TNBC breast tumors. In TNBC, mutant p53 is not a maker of the cancer stem cell (CSC) phenotype (*44*). Mutant p53 is a predictor of endocrine resistance in luminal B breast cancers, which is associated with CSC populations (*8, 45*). However, recent studies have associated mutant p53 with an altered transcriptional program leading to a stem cell phenotype in HER2-enriched breast cancers (*46*). A similar phenomenon likely occurs in TNBC and luminal B tumors; our transcriptomic analysis is the first evidence to support a role for mutant p53 and stem like features in these subtypes. Stem cell like tumors in our model are comprised of TNBC, HER2-enriched and luminal B tumors, and have the shortest survival amongst the tumor groups, suggesting mutant p53 drives a common transcriptome reflective of aggressive stem like features.

### Group B MaP^R245W/+^ Mammary Tumors Activate a Well-Differentiated & Enhanced Metabolic Active Program

Group B well-differentiated metabolically active tumors have two histological classifications: carcinomas with adenosis or papillary pattern (Fig. 2I, Table 1). These are mammary tumors that have retained their ductal structure and either have an increase in ducts and glands (adenosis pattern) or ducts with finger-like projections, giving them a well-differentiated appearance. Molecular subtyping of Group B *MaP^R245W/+^* mammary tumors revealed this group consists of 54% luminal A and 46% TNBC (Fig. S3B, Table 1). The TNBC tumors of this group likely represent the luminal TNBC subtype, which is driven by androgen receptor (*Ar)* (*47, 48*).

Comparison of Group B (n=13) mammary tumors to Groups A and C (n=38) revealed 3,192 DEGs, of which 988 genes are upregulated, and 2,204 genes are downregulated (Figs. S3I and J). GSEA analysis of Group B mammary tumors revealed significant pathway and process enrichments involving: KEGG Drug Metabolism Cytochrome P450, GOCC Apicolateral Plasma Membrane, KEGG Valine, Leucine and Isoleucine Degradation, Ctrl vs BLIMP1 Transduced GC B cell DN; Reactome Peroxisomal Lipid Metabolism, GOMF Structural Constituent of Muscle; Hallmark Estrogen Response Early, Reactome Keratinization; Tumoricidal vs Control Macrophage Up; and Hallmark Estrogen Response Late (Fig. 2F). Group B tumors activate a transcriptional program involved in altered bioenergetics and metabolism, estrogen signaling, and keratinization, characteristics of aggressive luminal breast tumors (*49–54*). Therefore, we classify Group B as Well-Differentiated Metabolically Active Tumors.

### Group C MaP^R245W/+^ Mammary Tumors Activate an Immunosuppressive Program

Group C consists of ten *MaP^R245W/+^* mammary tumors. These tumors are histologically classified as: metaplastic carcinoma with spindle cells, small blue cell tumor, sarcomatoid carcinoma with sarcoma-like and metaplastic features, and carcinoma (Fig. 2J, Table 1). All tumors within Group C, lack expression of *Esr1*, *Pgr*, and *Erbb2* (as measured by qRT-PCR), and are TNBC (Fig. S3C, Table 1).

Group C (n=10) mammary tumors compared to Groups A and B mammary tumors (n=41) revealed total of 4,338 DEGs, with 2,111 genes upregulated and 2,227 genes downregulated (Figs. S3K and L). GSEA revealed significant enrichment of the following pathways and processes: Adenosine A3R vs T Cell Membranes in Mast Cell Up; TLR1 TLR2 Ligand vs NOD and TLR1 TLR2 Ligand 24h Treated Monocyte up, Naïve CD8 T cell vs Monocyte Dn; Patel Skin of Body Zostavax Age 70 93YO Vsq Challenge 3 dy Up; WT vs MYD88 KO Macrophage 48hr MBOVIS BCG Stim Dn; B Cell vs Monocyte Dn; Lupus B Cell vs Lupus Myeloid Dn; Naïve CD4 T cell vs Monocyte Dn; Fletcher PBMC BCG 10W Infant BCG Stimulated vs Unstimulated 10W Dn (Fig. 2G). Group C mammary tumors did not activate any hallmark cancer pathways. Interestingly, these tumors activated processes involving adenosine signaling and genes upregulated in myeloid derived cells that mediate immune suppression, particularly monocytes. Enhanced adenosine signaling via the adenosine A3R receptor expressed on monocytes and other myeloid cells mediate immunosuppression (*55*). Specifically, activation of the A3R receptor leads to dampened T cell cytokine production and polarization of M1 macrophages to M2 macrophages, like Group C signatures associated with genes downregulated in naïve T cells and genes upregulated by macrophages (*55, 56*). Furthermore, elevated adenosine signaling is associated with enhanced activity of monocyte-myeloid derived suppressor cells (M-MDSCs) (*57*). The enhanced enrichment of immunosuppressive processes typically mediated by monocytes in Group C tumors, suggests these are very divergent from the epithelial cells of Groups A and B and thus, we will refer to Group C tumors as Immunosuppressive.

### MaP^R245W/+^ Mammary Tumors Harbor Group Specific Mutation Burdens and SNP Classifications

A total of 48 mammary tumors and matched normal tails and/or ear tissues were subjected to whole exome sequencing (WES). We identified a total of 6,992 nonsynonymous mutations across all tumors (Fig. 3A, Data S1-3). Missense mutations were the most common variant classification and single nucleotide polymorphism (SNP) was the most common variant across Groups A, B, and C mammary tumors (Figs. 3A and B). The median number of variants for Group A and B were 51 and 50, respectively, while the median number of variants for Group C was 236 (Fig. 3C). Group C tumors had the highest tumor mutation burden (TMB) equivalent to 3.6 per Mb compared to 1.3 per Mb for both Groups A and B mammary tumors (Fig. 3D). Differences in TMB across Groups A, B, and C were statistically significant (p-value < 0.0001, ANOVA) (Fig. 3E). Group specific analyses revealed that Group C mammary tumors diverge from the C>T SNP pattern of Group A and B tumors and distinctly harbor SNPs that were mostly T>G and T>C mutations, the majority of which were transversions (Fig. 3F). Enrichment of transversions has been associated with high TMB in human tumors; and Group C mammary tumors follow this phenomenon (*58*).

**Fig. 3.**
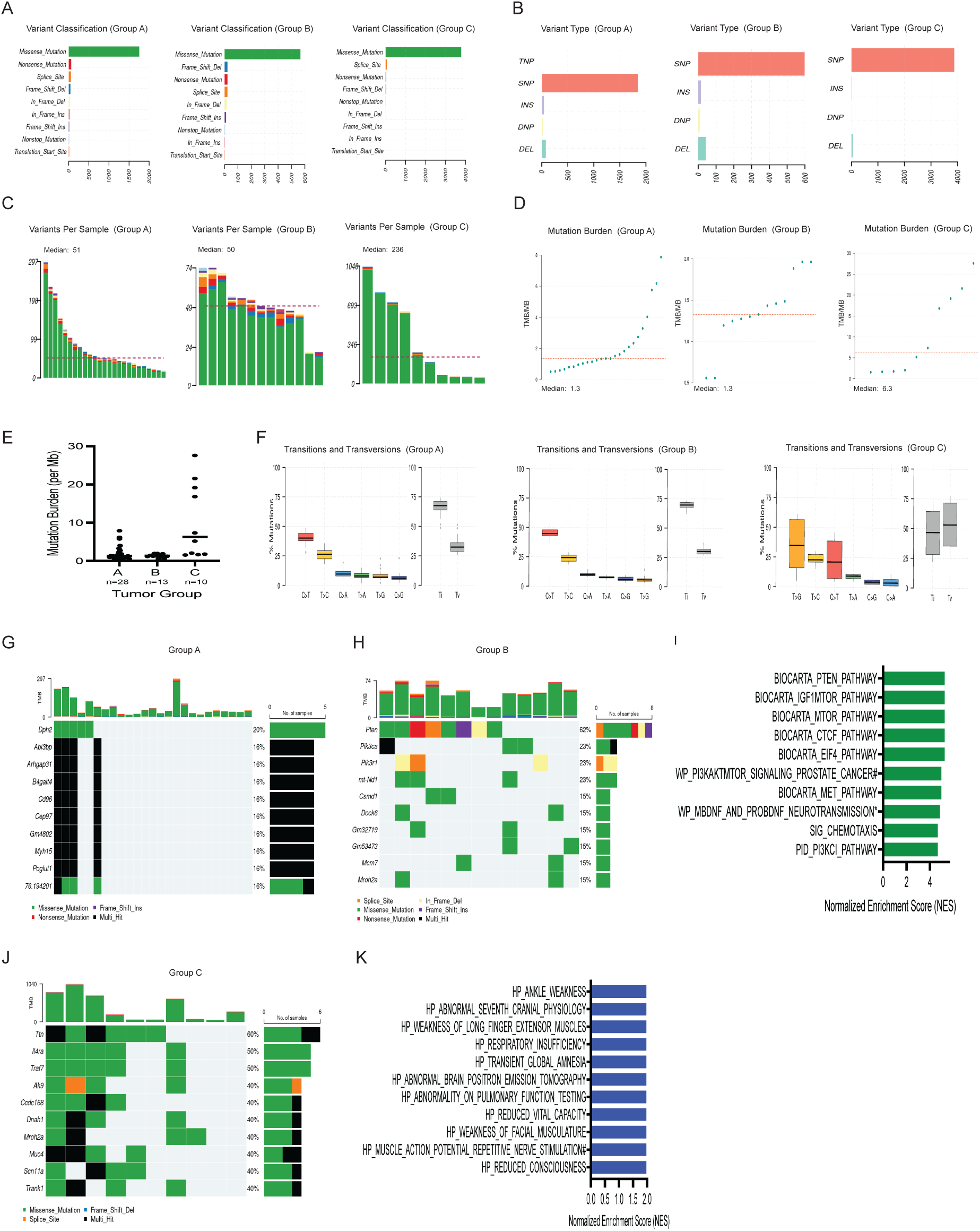
Mutational Landscape of *MaP^R245W/+^* Mammary Tumors. (A)Summary of Group Specific Variant Classification: Missense Mutation (green), Nonsense mutation (red), Splice Site (orange) Frame shift deletion (blue), In frame deletion (Yellow), In frame insertion (pink), Frame shift insertion (lavender), Nonstop mutation (light blue), Translation start site (peach) (B) Summary of Group Specific Variant Classes: TNP, SNP (peach), INS (lavender), DNP (Yellow); DEL (Teal) (C) Group Specific Summaries of Total Nonsynonymous Variants Per Sample (D) Group Specific Summaries of Tumor Mutation Burden (E) One way ANOVA Analysis Comparing Differences in Tumor Mutation Burden Across *MaP^R245W/+^* Tumor Subgroups (F) Group Specific Summaries of Transitions and Transversions. (G) Oncoplot of Recurrent mutations in Group A tumors. (H) Oncoplot of Recurrent mutations in Group B tumors. (I) Summary of GSEA enrichments Associated with Group B Top Ten Mutations Oncoplot of Recurrent mutations in Group C tumors. (J) Oncoplot of Recurrent Mutations in Group C Tumors. (K) Summary of GSEA enrichments Associated with Group C Top Ten Mutations Oncoplot of Recurrent mutations in Group C tumors.

### Loss of Heterozygosity Does Not Correlate with Group Specific Mutation Features

We sought to assess if loss of heterozygosity (LOH) was associated with the observed group specific differences in TMB and SNP classifications across *MaP^R245W/+^* mammary tumors. The status of the wild type (WT) *Trp53* allele was inferred from the WES data using the Control-FREEC algorithm. The B-allele frequency (BAF) was calculated at the C733T SNP to detect gains or losses with the *Trp53* locus. Across Group A tumors, 19 tumors retained heterozygosity, while 6 tumors underwent LOH (Table S2, Fig. S4). All Group B and C tumors retained heterozygosity at the *Trp53* locus as measured by Control-FREEC (Table S2, Fig. S5 and S6).

We also performed Sanger sequencing across 25 tumors representing Groups A (n=10), B (n=5), and C (n=10) to measure *Trp53* wild type allele retention. For Sanger sequencing, the retention of the WT allele was measured based on the ratio of peak amplitudes of wild type bases to mutant bases, as previously described (*22*). The Sanger sequencing data corroborated the FREE-C analysis, except for one outlier (Table S2, Figs. S7B-D). These analyses revealed no correlation between TMB load in Group C and LOH status. Collectively, 98% of *MaP^R245W/+^* mammary tumors retain heterozygosity at the *Trp53* locus (Table S2), indicating there is no pressure for these tumors to lose the wildtype allele.

### MaP^R245W/+^ Mammary Tumors Harbor Group Specific Recurrent Mutations

To identify group specific recurrent mutations, in an unbiased manner, mutation calling was performed for tumors in Groups A, B and C. The top 10 most recurrently mutated genes in Group A stem cell like mammary tumors were *Dph2* (5 tumors, 20%)*, Abi3bp* (4 tumors, 16%)*, Arhgap31* (4 tumors, 16%)*, B4galt4* (4 tumors, 16%)*, Cd96* (4 tumors, 16%)*, Cep97* (4 tumors, 16%)*, Gm4802* (murine specific pseudogene) (4 tumors, 16%)*, Myh15* (4 tumors, 16%)*, Poglut1* (4 tumors, 16%)*, and SRR518976.194201* (murine specific pseudogene) (4 tumors, 16%) (Fig. 3G). There were no cancer-related signaling pathways dysregulated among the top 10 mutated genes. However, GSEA analysis showed 5 out of 10 genes (*Cd96*, *Poglut1*, *Arhgap31*, *B4galt4*, and *Myh15*) correspond to the human cytogenetic band chr3q13, a chromosomal locus comprised of cooperatively acting tumor suppressor genes, which have been associated with loss of heterozygosity (LOH) and copy number alterations (CNAs) in various cancers (*59, 60*). Group A recurrent mutations were queried against the COSMIC database to assess if any mutations correspond to Tier 1 or Tier 2 known cancer-causing genes (*61, 62*). There were no recurrent Group A mutations identified as known cancer-causing genes. Group A recurrent mutations were observed in only 6 of 25 (24%) tumors (Fig. 3G). Therefore, Group A stem cell like mammary tumors do not have recurring mutations.

In the well-differentiated metabolically active Group B tumors, the 10 most recurrently mutated genes included *Pten* (8 tumors, 62%), *Pik3ca* (3 tumors, 23%), *Pik3r1* (3 tumors, 23%), *mt-Nd1* (3 tumors, 23%), *Csmd1* (2 tumors, 15%), *Dock6* (2 tumors, 15%), *Gm32719* (predicted murine specific gene) (2 tumors, 15%), *Gm5473* (predicted murine specific gene) (2 tumors, 15%), *Mcm7* (2 tumors, 15%), and *Mroh2a* (2 tumors, 15%) (Fig. 3H). GSEA analysis of the recurrent mutations (excluding predicted murine specific genes *Gm53473* and *Gm32719*), revealed dysregulation of the Pi3k/Akt/mTOR signaling pathway (Fig. 3I). The most significantly mutated gene in Group B tumors is the tumor suppressor *Pten*, which when mutated or deleted results in the activation of the Pi3k/Akt/mTOR pathway (PAM). Additional dysregulated pathways including IGF, CTCF, MET, and mBDNF/Pro BDNF converge upon the PAM pathway leading to its activation (*63–66*). The EIF4 pathway is downstream of the PAM pathway (*67*). Interestingly, each of the pathways identified in this enrichment analysis are attributed to mutations in *Pten*, *Pik3ca*, and *Pik3r1*. Eleven of thirteen (85%) Group B tumors harbor mutations in one or more of these three genes.

The two tumors lacking mutations in *Pten*, *Pik3ca*, and *Pik3r1* harbor mutations in one or more of the following genes: *Dock6*, *Gm53473, Mcm7,* and *Mroh2a*. Each of these genes are associated with PAM signaling. *Dock6* encodes a guanine exchange factor, that activates Rac1, leading to enhanced PAM signaling (*68*). *Gm53473* encodes a predicted murine gene of unknown function that is located within the plasma non-HDL cholesterol 3 (Pnhdlc3) expression quantitative trait locus (eQTL), and genes mutated at this locus are associated with abnormal non-HDL lipid levels. Oxidation of non-HDL lipids can fuel activation of the PAM signaling cascade in tumor cells (*69*). Mcm7 has been shown to have positive and negative regulation of Pi3k signaling in a context dependent manner in various cancers (*70, 71*). *Mroh2a* encodes a HEAT domain containing protein of unknown function. HEAT domains were originally identified in four proteins from which the domain name was derived: <u>h</u>untingtin, eukaryotic translation <u>e</u>longation factor 3, the regulatory subunit <u>A</u> of protein phosphatase 2A (PP2A), and mechanistic target of rapamycin (mTOR) (*72*). HEAT domain proteins are involved in intracellular signaling including regulation of the PAM pathway (*72*).

Other genes mutated in Group B mammary tumors include *Gm32719*, *mt-Nd1*, and *Csmd1*. *Gm32719* is a predicted gene of unknown function. *mt-Nd1*, encodes mitochondrial NADH dehydrogenase 1, which plays a critical role in ATP synthesis, and when mutated leads to increased production of reactive oxygen species (ROS). While mt-Nd1, is not directly associated with PAM signaling, elevated ROS leads to activation of this pathway (*73*). Lastly, *Csmd1*, encodes a protein involved in mammary gland development and when mutated results in upregulation of Pi3k and its downstream signaling (*74*). Therefore, Group B tumors are universally defined by one or more mutations that enhance PAM signaling.

Each of the recurrently mutated genes identified in Group B tumors were queried in the COSMIC database. *Pten*, *Pik3ca,* and *Pik3r1* were identified as Tier 1 or Tier 2 known cancer-causing genes (Table 2). The ConVarT Analysis tool was used to identify human counterparts to the cancer-causing murine variants and if any conserved variants are pathogenic (*75*). There were 8 different *Pten* variants identified in our WES analysis. Except for a splice site variant, all murine *Pten* variants were highly conserved in humans and associated with pathogenicity (Table 2). There were 4 *Pik3ca* variants in Group B tumors, each of which were also highly conserved and pathogenic. Three *Pik3r1* variants were identified. One variant was a splice site mutation and occurred in a conserved region for a human pathogenic variant. Two in-frame deletions occurred, one of which corresponds to human variants of unknown significance and the other corresponds to a human pathogenic variant (Table 2). In many cancers, *PTEN* and *PIK3CA* mutations are mutually exclusive to *TP53* mutations (*76*). However, co-mutation of *TP53* with *PIK3CA* or *PTEN* leads to aggressive disease (*77, 78*). Thus, Group B tumors are defined by alterations in established cancer-causing genes that activate the PAM pathway.

**Table 2.**
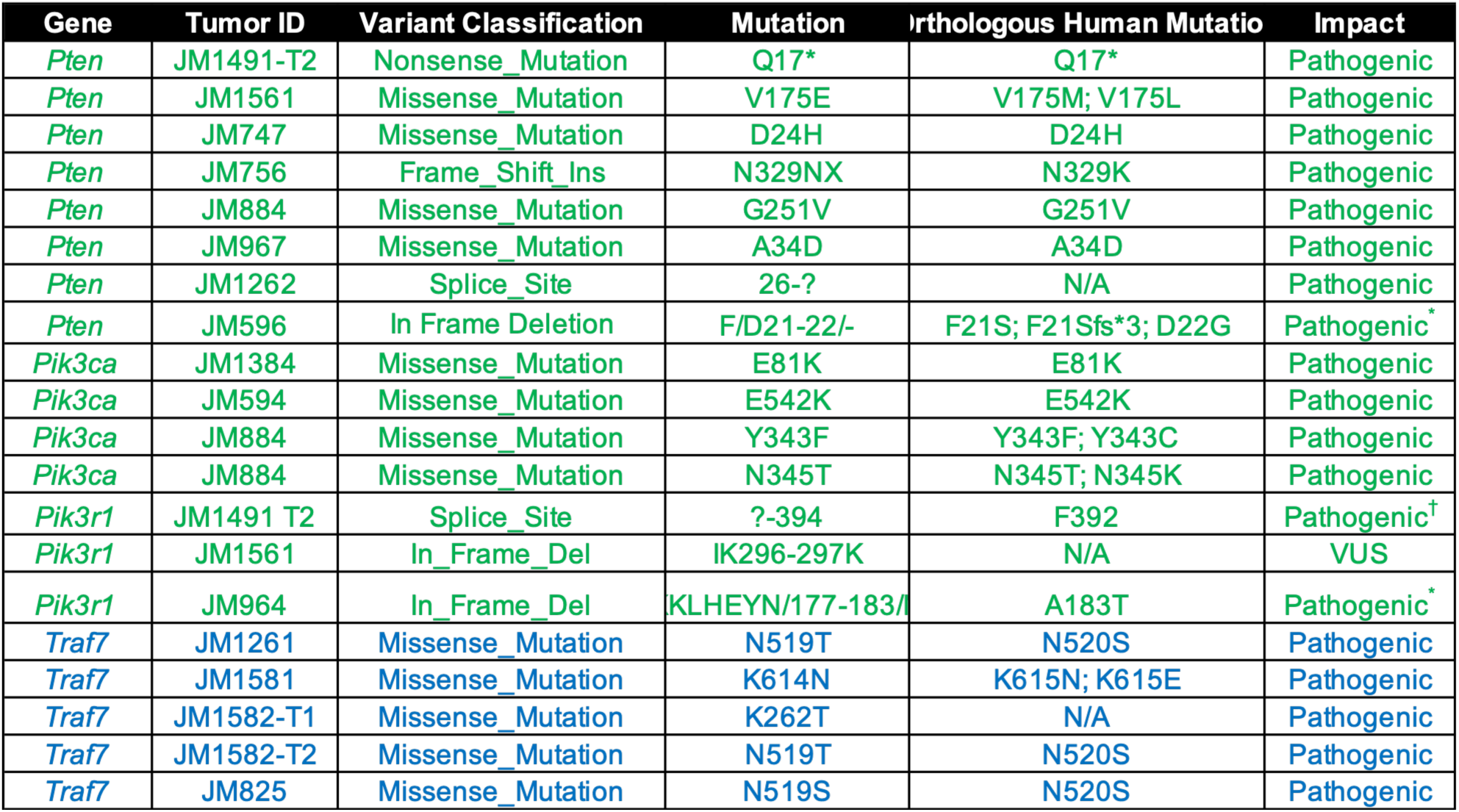
Group Specific Pathogenic Mutations in *MaP^R245W/+^* Mammary Tumors. Summary of group specific recurrent mutations in COSMIC-designated cancer-causing genes. Green: Group B Well-Differentiated Metabolic Active Tumors. Blue: Group C Immunosuppressed Tumors. *Pathogenic mutation near orthologous SNP variant in human. †Pathogenic mutation near orthologous splice site variant in human. VUS: Variant of Unknown Significance.

| Gene | Tumor ID | Variant Classification | Mutation | Orthologous Human Mutatio | Impact |
| --- | --- | --- | --- | --- | --- |
| *Pten* | JM1491-T2 | Nonsense_Mutation | Q17* | Q17* | Pathogenic |
| *Pten* | JM1561 | Missense_Mutation | V175E | V175M; V175L | Pathogenic |
| *Pten* | JM747 | Missense_Mutation | D24H | D24H | Pathogenic |
| *Pten* | JM756 | Frame_Shift_Ins | N329NX | N329K | Pathogenic |
| *Pten* | JM884 | Missense_Mutation | G251V | G251V | Pathogenic |
| *Pten* | JM967 | Missense_Mutation | A34D | A34D | Pathogenic |
| *Pten* | JM1262 | Splice_Site | 26-? | N/A | Pathogenic |
| *Pten* | JM596 | In Frame Deletion | F/D21-22/- | F21S; F21Sfs*3; D22G | Pathogenic* |
| *Pik3ca* | JM1384 | Missense_Mutation | E81K | E81K | Pathogenic |
| *Pik3ca* | JM594 | Missense_Mutation | E542K | E542K | Pathogenic |
| *Pik3ca* | JM884 | Missense_Mutation | Y343F | Y343F; Y343C | Pathogenic |
| *Pik3ca* | JM884 | Missense_Mutation | N345T | N345T; N345K | Pathogenic |
| *Pik3r1* | JM1491 T2 | Splice_Site | ?-394 | F392 | Pathogenic† |
| *Pik3r1* | JM1561 | In_Frame_Del | IK296-297K | N/A | VUS |
| *Pik3r1* | JM964 | In_Frame_Del | KLHEYN/177-183/ | A183T | Pathogenic* |
| *Traf7* | JM1261 | Missense_Mutation | N519T | N520S | Pathogenic |
| *Traf7* | JM1581 | Missense_Mutation | K614N | K615N; K615E | Pathogenic |
| *Traf7* | JM1582-T1 | Missense_Mutation | K262T | N/A | Pathogenic |
| *Traf7* | JM1582-T2 | Missense_Mutation | N519T | N520S | Pathogenic |
| *Traf7* | JM825 | Missense_Mutation | N519S | N520S | Pathogenic |

The ten most recurrently mutated genes in Group C immunosuppressive tumors include *Ttn* (6 tumors, 60%), *Il4ra* (4 tumors, 40%) (orthologous to human *IL4R*), *Traf7* (4 tumors, 40%), *Ak9* (4 tumors, 40%), *Ccdc168* (4 tumors, 40%), *Dnah1* (4 tumors, 40%), *Mroh2a* (4 tumors, 40%), *Muc4* (4 tumors, 40%), *Scn11a* (4 tumors 40%) and *Trank 1* (4 tumors, 40%) (Fig. 3J). GSEA analysis of the recurrently mutated genes revealed dysregulation of human physiological processes related to muscle, central nervous system, and pulmonary function (Fig. 3K). These dysregulated physiological processes are largely due to *Ttn*, *Ak9*, *Traf7*, and *Dnah1* mutations.

While the recurrently mutated genes do not participate in a common pathway, most of the mutated genes have associations with high TMB and/or immune regulation in other cancer contexts. *TTN*, encodes the protein titin, the largest protein in the human genome and plays a critical role in the structure and function of striated muscle. Titin’s long coding sequence allows it to accumulate a high number of mutations and is associated with a high TMB and enhanced immunogenicity in various cancers (*79, 80*). In various cancers, *TTN* mutations are associated with response to immunotherapy and longer survival (*79–81*). Thus, *Ttn* mutations may contribute to the high TMB and longer survival exhibited by the Group C mice. *Il4ra* is the murine homolog to *IL4R*, which encodes interleukin-4 receptor alpha, which bind IL4 and IL13 to mediate immune suppression. In cancer, mutations in *IL4R* lead to strengthened immunosuppression via M2 macrophage expansion, recruitment of regulatory T cells (T regs); and inhibiting the function of cytolytic T cells (*82–84*). Furthermore, *IL4R* mutations correlate with immunotherapy resistance in certain tumor types (*85*).

*Ak9* encodes adenylate kinase 9, which catalyzes key reactions involved in adenine homeostasis, resulting in adenosine imbalance, a contributor of immune suppression. (Klepinin 2020) Adenosine signaling is enriched in the transcriptomes of our immunosuppressive tumors and may be attributed to *Ak9* mutations. AK9 has tumor suppressive functions in lung cancer, and it is thought that mutations in this gene lead to deregulated nucleotide bioenergetics and cancer progression (*86*). Thus, Ak9 may be a tumor suppressor in Group C tumors and may contribute to immune suppression in this group. *Ccdc168*, encodes coiled-coiled domain containing-168, which when downregulated is associated with enhanced immunotherapy response in prostate cancer (*87*). *Muc4*, encodes mucin 4, a cell surface glycoprotein. Co-mutations in *MUC4* and *TTN* are associated with high mutation burden and positive response to immunotherapy in gastric cancers and other cancers (*88*). *Muc4* and *Ttn* mutations may cooperate in Group C to promote an environment favorable to immunotherapy response.

Cosmic analysis revealed Group C tumors recurrently mutate a single Tier 1 cancer causing gene: *Traf7* (Table 2). TRAF7, TNF receptor-associated factor 7, is a signaling adaptor of multiple inflammatory pathways in muscle and brain (*89*). Four *Traf7* variants were identified in Group C mammary tumors. Except for one variant, all *Traf7* variants had a highly conserved human counterpart associated with pathogenicity (Table 2). TRAF7 has a role in development and progression in cancer (*90, 91*). In breast cancer, TRAF7 promotes ubiquitin-proteosome mediated degradation of p53 (*92*). Mutations in *TRAF7* have been associated with mesenchymal tumors that are highly aggressive (*89*). Similarly, 80% of Group C immunosuppressive tumors are comprised of rare aggressive tumors known to have mesenchymal characteristics including metaplastic carcinoma with spindle cells, small blue cell tumor and sarcomatoid carcinoma with metaplastic features (*93, 94*) (Fig. 2J and Table 1).

Collectively, mutational profiling suggests *MaP^R245W/+^* tumors have distinct intrinsic dysregulated pathways and processes. Group B mammary tumors likely activate the PAM pathway via *Pten, Pik3ca, and Pik3r1* mutations. Group C mammary tumors preferentially mutate muscular, CNS and pulmonary genes with varied roles in modulating the immune system.

### CNA Analysis Reveals Potential Group Specific Mediators of Metastatic Progression

Somatic copy number alterations (CNA) were inferred from WES data of the 48 *MaP^R245W/+^* mammary tumors (Fig. S8). Group specific recurrent CNAs were identified using the GISTIC 2.0 algorithm. Recurrent CNA analysis performed across 25 Group A mammary tumors revealed 9 significant amplifications and 16 deletions (q value <0.25) (Figs. 4A-C and Data S4). In contrast to their lack of recurrent mutations, Group A tumors have high copy number alterations. Biologically relevant frequent copy number gains occurred in chromosomes 2q, 6q, 7q, 8q, 9q, 11q, 13q, 16q, and 17q (Figs. 4A and C and Table S4). Frequent losses with potential biological relevance occurred in chromosomes 4q, 5q, 6q, 7q, 8q, 9q, 10q, 11q, 12q, 13q, 16q, 17q, and 18q (Figs. 4B and C and Data S4). All recurring CNAs were queried against the COSMIC database to identify gains and losses of known oncogenes and tumor suppressors. Recurring amplifications in oncogenes included *Met* (6qA2, 44%); *Birc3*, *Yap1* (9qA1, 40%); and *Fgfr4* and *Nsd1* (13qB1, 40%) (Figs. 4A and C, Table 3 and Data S4). Recurrent deletions in tumor suppressors included *Nf1* (11qB5, 36%), *Pik3r1* and *Rad17* (13qD2.1, 24%) (Figs. 4B and C, Table 3 and Data S4).

**Fig. 4.**
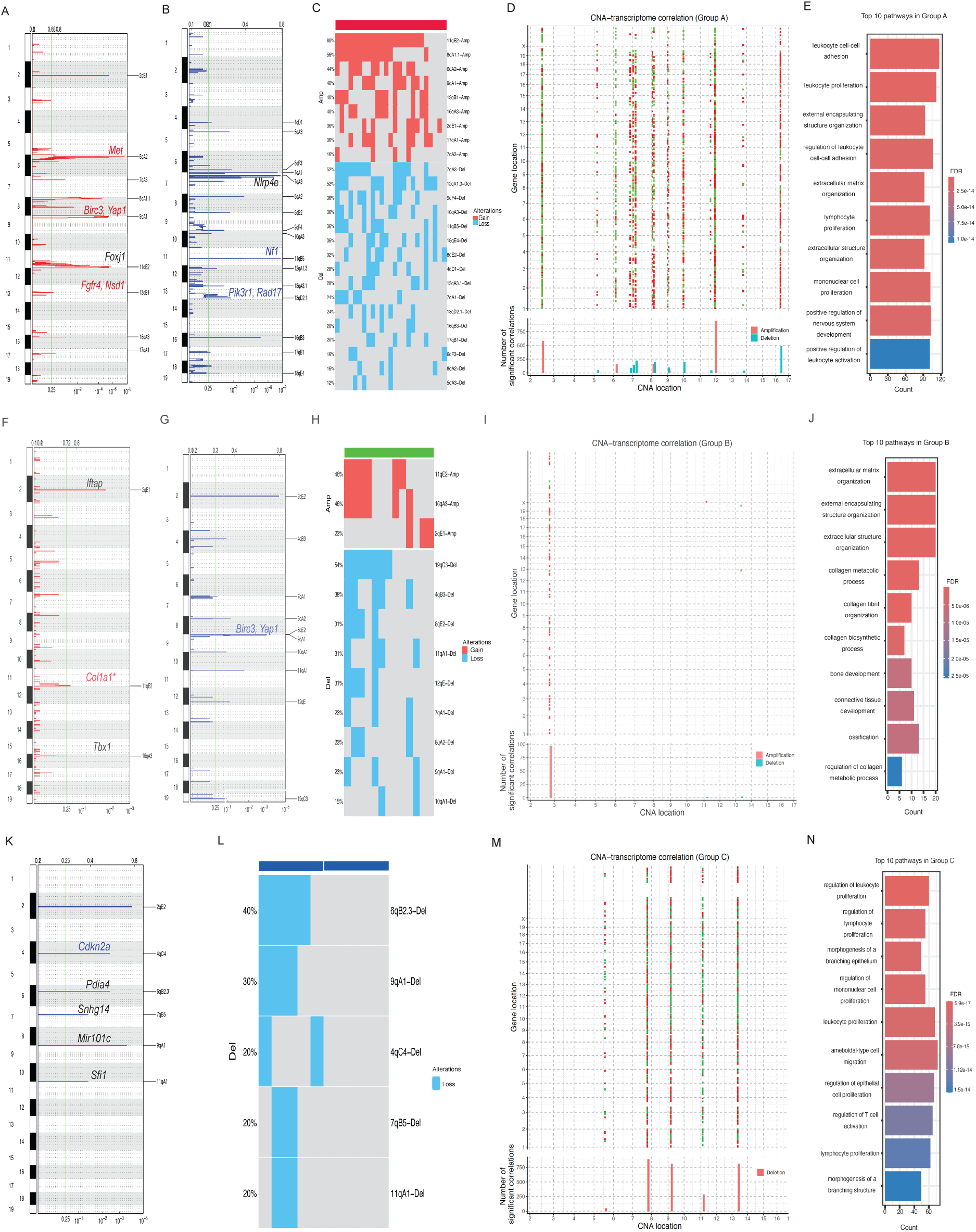
Genomic Landscape of *MaP^R245W/+^* Mammary Tumors. (A) Genome-wide recurring focal amplifications of Group A tumors with GISTIC 2.0 FDR q-values on the bottom. Significant peaks were annotated with COSMIC designated driver oncogenes in red. (B) Genome-wide recurrent focal deletions of Group A tumors with GISTIC 2.0 FDR q-values on the bottom. COSMIC designated driver tumor and/or metastasis suppressors within deletion peaks are annotated in blue. Genes involved in CNA-mRNA regulation annotated in black. (C) Frequency plot of significant amplifications and deletions in Group A mammary tumors. (D) Summary of copy number alterations significantly correlated with differentially expressed genes in Group A mammary tumors. (E) Biological and molecular processes significantly enriched in differentially expressed genes correlated with significant copy number alterations in Group A mammary tumors. (F) Genome-wide recurring focal amplifications of Group B tumors with GISTIC 2.0 FDR q-values on the bottom. Significant peaks were annotated with COSMIC designated driver oncogenes in red. *indicates a locus harboring more than two COSMIC cancer-causing genes. Genes involved in CNA-mRNA regulation annotated in black. (G) Genome-wide recurrent focal deletions of Group B tumors with GISTIC 2.0 FDR q-values on the bottom. COSMIC designated driver tumor and/or metastasis suppressors within deletion peaks are annotated in blue. (H) Frequency plot of significant amplifications and deletions in Group B mammary tumors. (I) Summary of copy number alterations significantly correlated with differentially expressed genes in Group B mammary tumors. (J) Biological and molecular processes significantly enriched in differentially expressed genes correlated with significant copy number alterations in Group B mammary tumors. (K) Genome-wide recurrent focal deletions of Group C tumors with GISTIC 2.0 FDR q-values on the bottom. COSMIC designated driver tumor and/or metastasis suppressors within deletion peaks are annotated in blue. Genes involved in CNA-mRNA regulation annotated in black. (L) Frequency plot of significant deletions in Group C mammary tumors. (M) Summary of copy number alterations significantly correlated with differentially expressed genes in Group C mammary tumors. (N) Biological and molecular processes significantly enriched in differentially expressed genes correlated with significant copy number alterations in Group C mammary tumors.

**Table 3.** Group Specific Pathogenic Copy Number Alterations in *MaP^R245W/+^* Mammary Tumors. Summary of group specific recurrent copy number alterations in COSMIC-designated cancer-causing genes.

| Murine CNA | Orthologous Human COSMIC Genes at Syntenic Locus | Group Membership |
| --- | --- | --- |
| 6qA2-Amp | *Met* | Group A |
| 9qA2-Amp | *Birc3, Yap1* | Group A |
| 13qB1-Amp | *Fgfr4, Nsd1* | Group A |
| 11qB5-Del | *Nf1* | Group A |
| 13qD2.1-Del | *Pik3r1, Rad17* | Group A |
| 11qE2-Amp | *Aspr1, Axin2, Brca1, Cant1, Ccr7, Cd79b, Cdk12, Col1a1, Ddx15, Erbb2, Etv4, H3f3b, Ikzf3, Kat7, Lasp1* | Group B |
| 9qA2-Del | *Birc3, Yap1* | Group B |
| 4qC4-Del | *Cdkn2a* | Group C |

Expression quantitative trait loci (eQTL) analysis was performed to identify *cis* and *trans* effects of Group A CNAs on differential gene expression, (Fig. 4D and Data S5). Specifically, differential gene expression analysis was performed on tumors harboring significant amplifications and deletions compared to those without them. *Cis*-regulatory events were defined as DEGs located on the same chromosome of a given CNA. *Trans-*regulatory events were defined as DEGs that were not located on the same chromosome as a CNA. There were 4,357 significant DEGs associated with a CNA (Fig. 4D and Data S5). There were 325 DEGs *cis*-regulated, and 4,032 DEGS *trans-*regulated by CNAs. There was no evidence of broad *cis-*regulatory effects of CNAs on DEGS. DEGS were predominantly enriched in biological and molecular processes involving leukocyte cell adhesion, proliferation and activation; extracellular matrix organization; nervous system development; and lymphocyte and mononuclear cell proliferation (Fig. 4E).

The most notable CNAs with *trans* effects were on chromosomes 2q, 5q, 6q,7q, 8q, 9q, 11q, 13q, 16q and 17q (Fig. 4D). Among them, 11q the most frequently amplified locus (80%), contains *Foxj1*, a transcriptional regulator of ciliogenesis which has been associated with the neural stem cell niche and metastasis in various cancers (Figs. 4A, C and D and Tables S4 and S5) (*95–97*). One of the most frequent deletions (52%) occurred on chromosome 7q, containing *Nlrp4e* (Figs. 4B, C and D and Tables S4 and S5). *Nlrp4e* is orthologous to human *NLRP4*, which encodes a protein that activates NK cells to mediate tumor suppressive functions (*98*). It is possible deletion of *Nlrp4e* is associated with the gene signature enrichment associated with NK cell inactivation observed in the Group A transcriptome (Fig. 2E).

GISTIC 2.0 analysis in the 13 Group B well-differentiated metabolically active tumors revealed 3 significant amplifications and 9 deletions (q value <0.25) (Figs. 4F-H and Data S6). Frequent copy number gains occurred in chromosomes 2q, 11q, and 16q (Figs. 4F and H and Data S6). Frequent losses with potential biological relevance occurred in chromosomes 4q, 7q, 8q, 9q, 10q, 11q, 12q, and 19q (Figs. 4G and H and Data S6). All recurring oncogenic amplifications, designated by COSMIC, occurred on the 11qE2 amplicon (23%) and included *Aspr1*, *Axin2*, *Brca1*, *Cant1*, *Ccr7*, *Cd79b*, *Cdk12*, *Col1a1*, *Ddx15*, *Erbb2, Etv4*, *H3f3b*, *Ikzf3*, *Kat7*, and *Lasp1* (Figs. 4F and H, Table 3 and Data S6). It is of interest to note many of the oncogenes in this amplicon have been associated with poor prognosis and/or metastasis in breast cancer. Recurrent deletions occurred in two COSMIC designated tumor suppressors, *Birc3 and Yap1 (*9qA1, 23%) (Figs. 4G and H, Table 3 and Data S6).

Similar eQTL analysis of Group B well-differentiated metabolically active tumors revealed 112 DEGS whose expression significantly correlates with CNAs (Fig. 4I and Data S7). There were 11 DEGs *cis*-regulated, and 101 DEGS *trans-*regulated by CNAs. There were no broad *cis-*regulatory effects of CNAs on DEGS. Most DEGS correlating with CNAs were predominantly enriched in processes involving extracellular matrix (ECM) organization, collagen organization and connective tissue development (Fig. 4J). These processes do not overlap with any of the dysregulated pathways identified in the transcriptomic analysis of Group B mammary tumors. Therefore, regulation of the ECM and connective tissue are distinctive processes *trans*-regulated by aberrant CNAs in Group B.

Loci with *trans* effects were on chromosomes 2q, 9q, 11q and 16q (Fig. 4I). Regarding amplifications, only two loci had *trans* effects on mRNA, 16qA3 (46%) and 2qE (23%) (Fig. 4I and Data S7). The 16qA3 locus contains *Tbx1*, which encodes the T box transcription factor, involved in cardio-pharyngeal development (Figs. 4F, H and I and Data S6 and S7) (*99*). Recent studies demonstrate that Tbx1 acts as an oncoprotein to promote cell cycle progression in breast cancer (*100*). The 2qE1 locus contains the murine predicted gene *B230118H07Rik*, which is orthologous to the human *IFTAP* gene (also referred to as *NWC*) (Figs. 4F, H, and I and Data S6 and S7). The *IFTAP* gene is the third gene part of the *RAG1/RAG2* locus and functions as a transcriptional repressor and has roles in the development of cilia, cell surface antenna that regulate the ECM (*101*). Deletions at 11q and 9q had minimal *trans* effects on gene expression, as we observed correlative differential expression of three genes across these two loci (Fig. 4I and Data S7). Therefore, amplification of *Tbx1* and *Iftap* likely drive the *trans* effects on the Group B transcriptome to regulate the ECM and connective tissue processes (Fig. 4J).

GISTIC 2.0 analysis was performed on 13 Group C immunosuppressed mammary tumors. Recurrent CNAs occurred in 5 deletions (q value < 0.25) (Figs. 4K and Data S8). Frequent copy number losses occurred at chromosomes 4q, 6q, 7q, 9q and 11q. (Figs. K and L and Supplementary Table 10). Recurrent deletions occurred in a single COSMIC defined tumor suppressor, *Cdkn2a* (4qC4, 20%) (Figs. 4K and L, Table 3, and Data S8).

eQTL analyses of Group C tumors revealed a total of 2,836 DEGs correlating with one deletion event (Fig. 4M and Data S9). The most significant biological processes affected include epithelial cell, leukocyte, lymphocyte and mononuclear cell proliferation; morphogenesis of a branching epithelium; ameboidal-type cell migration; and regulation of T cell activation (Fig. 4N). Regulation of branching morphogenesis and ameboidal-type cell migration potentially reflect a metastatic mechanism distinct from Groups A and B tumors.

Significant deletion events with *trans* effects on the transcriptome included 4q, 6q, 7q, 9q, and 11q (Fig. 4M and Data S9). The most frequently deleted locus in Group C mammary tumors was at 6qB2.3 (40%) and has significant *trans* effects on mRNA (Figs. 4K, L and M and Data S8 and S9). This locus contains the gene *Pdia4,* which functions in protein folding and endoplasmic reticulum homeostasis (Data S8 and S9) (*102*). Recent studies demonstrate a tumor suppressor role for Pdia4 in lung adenocarcinomas (*102*). The 9qA1 locus (30%) contains the putative tumor suppressor *Mir101c*, which suppresses proliferation, invasion and stem cell like phenotypes and activates apoptosis in various cancers (Fig. 4K and L and Data S8 and S9) (*103*).

The 4qC4 locus (20%) contains the COSMIC designated tumor suppressor *Cdkn2a* which is involved in regulation of the cell cycle (Table 3). Deletion of *Cdkn2a* is correlated with reduced T cell infiltration in melanoma and may contribute to the immunosuppressive phenotype of Group C tumors mammary tumors (*104*). The 7qB5 locus (20%) contains *Snhg14* (Figs. 4K and L and Data S8), a long non-coding RNA that regulates other small RNAs, leading to indirect regulation of genes involved in proliferation and survival (*105*). Lastly, the 11qA1 locus (20%) harbors the *Sfi1* gene (also referred to as *Pisd*) (Figs. 4K and L and Data S8), which encodes a protein involved in the structural dynamics of centrosome-associated contractile fibers. It is thought to have some tumor suppressive function, as its downregulation is associated with poor prognosis in breast cancer (*106*). Group C tumors harbor few CNAs; however, they preferentially delete the well-established tumor suppressor Cdk2na to regulate branching morphogenesis and immune cell proliferation and activation (Fig. 4N).

### MaP^R245W/+^ Mammary Tumors Have Distinct Immune Microenvironments That Mediate Immune Suppression

Given that the *MaP^R245W/^*^+^ mammary tumors have intrinsic heterogeneity profiles at both the transcriptomic and genomic levels, we hypothesized each intrinsic subgroup will have distinct immune microenvironments. Utilizing our bulk RNA-sequencing data, we implemented the CIBERSORT program (*107*) to infer which immune cell-types are present in each tumor group (Fig. 5A). Specifically, Group A stem cell like tumors were significantly enriched for resting dendritic cells (p=4.7 x 10^-05^) compared to Groups B and C mammary tumors (Fig. 5A). Resting dendritic cells function to mediate immune tolerance by sending negative signals to CD8^+^ T cells, via immune checkpoint molecules such as PD-1 and CTLA-4, resulting in T cell tolerance (*108*). No other immune cell types were significantly enriched in Group A stem cell like tumors. In contrast to the immunotolerant environment of Group A, Group B well-differentiated metabolically active tumors are enriched for two immune cell types: activated dendritic cells (p=0.00561), and CD8^+^ T cells (p=0.02837) (Fig. 5A). Activated dendritic cells present antigens to CD8^+^ T cells to stimulate adaptive immunity (*109*). Therefore, Group B well-differentiated metabolically active tumors represent a warmer immune microenvironment, compared to Groups A and C tumors, and may be more susceptible to immune checkpoint inhibitors. Lastly, immune cell type proportionate estimates for Group C immunosuppressive tumors had the highest enrichment of M2 macrophages (p=0.00055), which are alternatively activated macrophages that function to inhibit T cell activity and mediate immune suppression (*110*), often resulting in metastatic progression.

**Fig. 5.**
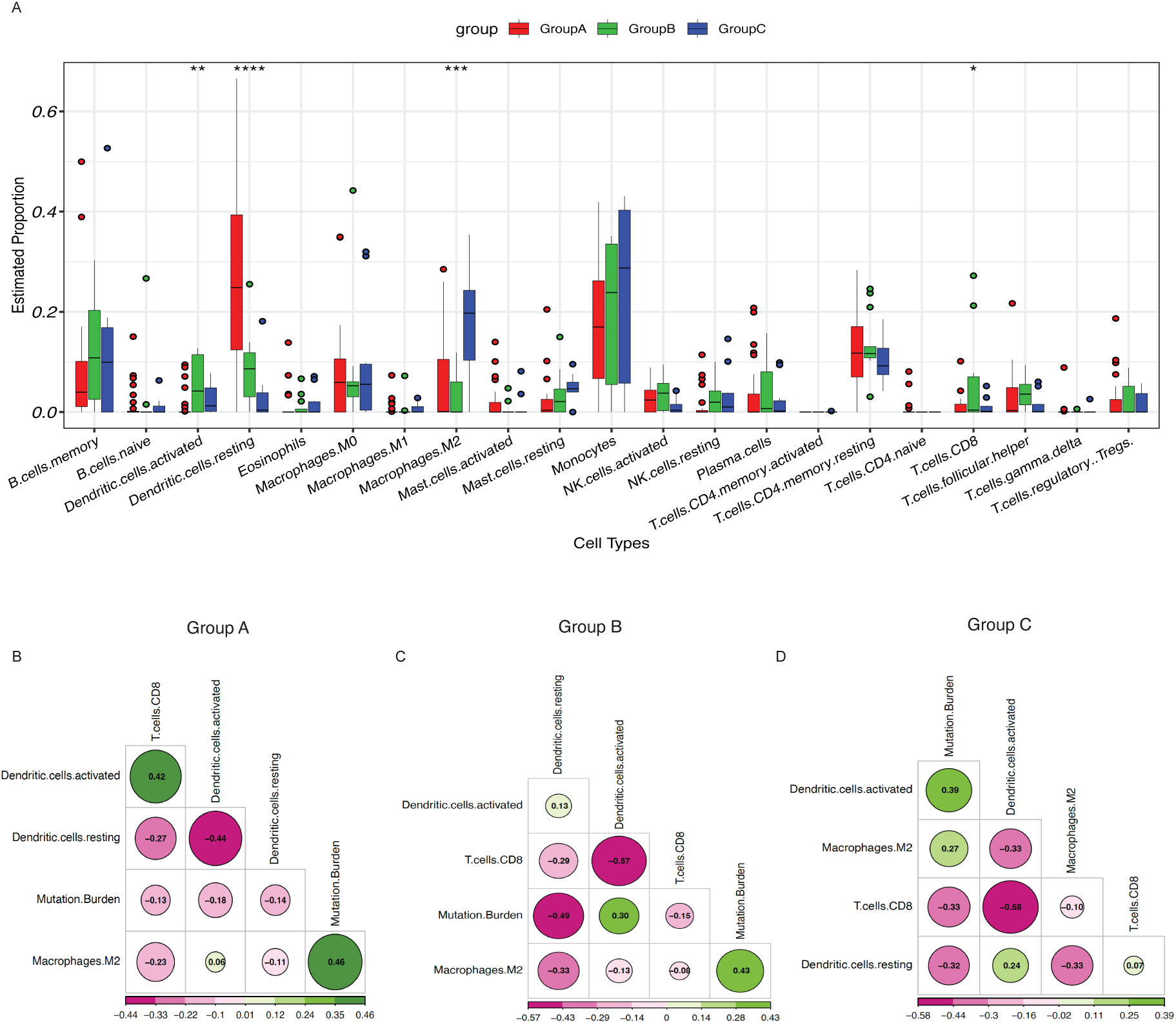
The Immune Landscape of *MaP^R245W/+^* Mammary Tumors. (A)CIBERSORT analysis to infer immune cell estimated proportions from bulk RNA-sequencing data of Groups A, B, and C mammary tumors. (B-D) Matrices summarizing group specific correlations between activated dendritic cells, resting dendritic cells, tumor mutation burden, and CD8 T cells in Groups A, B and C mammary tumors, respectively.

We next sought to determine if there are correlations among the group specific immune cell estimates and if there is correlation of the immune cell estimates with tumor mutation burden. In Group A mammary stem cell like tumors, we observed a significant positive correlation between TMB and M2 macrophages (r=0.46 p-value <0.05) (Fig. 5B). We observed a significant negative correlation between resting dendritic cells and activated dendritic cells (r= -0.44, p-value <0.05) (Fig. 5B). Group A tumors also had a significant positive correlation between CD8^+^ T cells and activated dendritic cells (r=0.42, p-value <0.05) (Fig. 5B). Overall, immune cell estimates suggest that even though Group A tumors can recruit dendritic cells into their tumor microenvironment (Fig. 5A), due to a lack of activating signals they are in turn inefficient at activating CD8^+^ T cells, corroborating the immune tolerance phenotype in this group. Group A tumors likely mediate immune suppression via inactive dendritic cells and M2 macrophages.

In Group B well-differentiated metabolically active tumors, which had the highest enrichment of CD8^+^ T cells; we identified a single significant negative correlation between CD8^+^ T cells and activated dendritic cells (r= -0.57, p-value <0.05) (Fig. 5C). Our CIBERSORT analysis revealed significant enrichment of both immune cell types (Fig. 5A), indicative of a warmer immune environment. Increased infiltration of cytotoxic CD8^+^ T cells leads to tumor control and improved outcomes in patients with breast cancer (*111, 112*). Plasticity in activation states of dendritic cells can potentiate opposing anti-tumor immunological programs (*113*). Activated dendritic cells can also possess more of a tolerogenic phenotype, leading to CD8^+^ T cell suppression, and an immune suppressed environment (*114*). Certain subsets of dendritic cells have also been shown to restrain their immunostimulatory function and downstream activation of T cells due to differential uptake of tumor antigens in an IL-4/IL-13 dependent manner (*115*). It is possible a similar phenomenon occurs in Group B tumors. Overall, Group B mammary tumors have a mixed state of infiltrating immune cells that include enrichment for cytotoxic CD8^+^ T cells.

Intergroup correlation analysis between group specific enriched immune cells and TMB in Group C immunosuppressive tumors did not reveal any significant correlations (Fig. 5D). Collectively, our CIBERSORT immune microenvironment estimates, and correlation analyses suggest *MaP^R245W/+^* tumors exhibit infiltrating immune cell heterogeneity, each associated with unique underlying immune suppression mechanisms.

### MaP^R245W/+^ Intrinsic Subtype of mBC are Recapitulated in Human Breast Tumors

Our -omics based analyses have allowed us to identify three novel intrinsic subtypes of breast cancer characterized by distinct transcriptomic and genomic profiles (Table 4). The stem-cell like tumors activate ribosome biosynthesis and E2F signaling (Table 4). These tumors harbor amplifications in *Met*, *Birc3*, *Yap1* and deletions in *Nf1* and *Rad17* (Table 4). Well-differentiated metabolically active tumors histologically retain ductal structures giving a well-differentiated appearance, activate estrogen and metabolic signaling, and have genomic alterations in *Pten* and/or *Pik3ca* (Table 4). Immunosuppressed tumors enrich for processes related to immune suppression (Table 4). We next sought to determine if our intrinsic subtypes of mBC are present in human breast tumors. Given that *TP53* mutations are associated with advanced breast cancer, we queried human breast tumors harboring a p53 missense mutation, from the TCGA-BRCA and METABRIC datasets for the presence of our intrinsic subgroups (*15, 116*).

**Table 4.** Group Specific Molecular Characteristics of *MaP^R245W/+^* Mammary Tumors. Summary of group specific pathways and processes, mutations, amplifications and deletions.

| Intrinsic Group | Enriched Pathways & Processes | Mutations | Amplifications | Deletions |
| --- | --- | --- | --- | --- |
| Stem Cell Like | Ribosome biosynthesis, E2F signaling | No recurrent | *Met, Birc3, Yap1* | *Nf1 , Rad17* |
| Well-Differentiated Metabolically Active | Cytochrome P450 metabolism, lipid metabolism, estrogen signaling | *Pten, Pik3ca, Pik3r1* | *Col1a1* * | *Birc3, Yap1* |
| Immunosuppressed | Immune suppression | *Traf7* | No recurrent | *Cdkn2a* |

Specifically, we performed a similarity cross species analysis, for which the expression levels of genes defining our murine transcriptomic subgroups from our original PCA analysis were queried in human tumors. Tumors were assigned a normalized similarity score to relate them to the A, B or C groups in our murine model. We identified 214 breast tumors within the TCGA-BRCA dataset that harbor a *p53* missense mutation. We included all *TP53* missense mutations in our analyses, as there are not sufficient *TP53^R248^* human samples in the TCGA and METABRIC cohorts to perform a variant specific analysis. Our similarity analysis revealed the presence of our three intrinsic subgroups in these breast tumors (Figs. 6A and B). Within the METABRIC dataset, 394 breast tumors had a *p53* missense mutation. Like the TCGA-BRCA dataset, our intrinsic gene signatures were present amongst the METABRIC tumors (Figs. 6C and D). In each analysis, dimension reduction PCA analysis does show some degree of overlap between groups in the human tumors. However, this may be due to differences between these human breast tumors and our murine tumors. For instance, it is unknown if *TP53* is the initiating mutation in human tumors. Secondly, most breast tumors in TCGA and METABRIC are stage 2/3 cancers. Our murine system has primary breast tumors and gross metastases at endpoint. Nonetheless, each of these analyses corroborated that our intrinsic signatures are not restricted to a specific molecular subtype of breast cancer (Figs. 6B and D). Importantly, these analyses revealed that these metastatic signatures are not exclusive to the p53R248 missense mutation. Therefore, the presence of a p53 missense mutation can lead to the activation of any one of three distinct intrinsic transcriptomic signatures associated with metastasis in breast cancer.

**Fig. 6.**
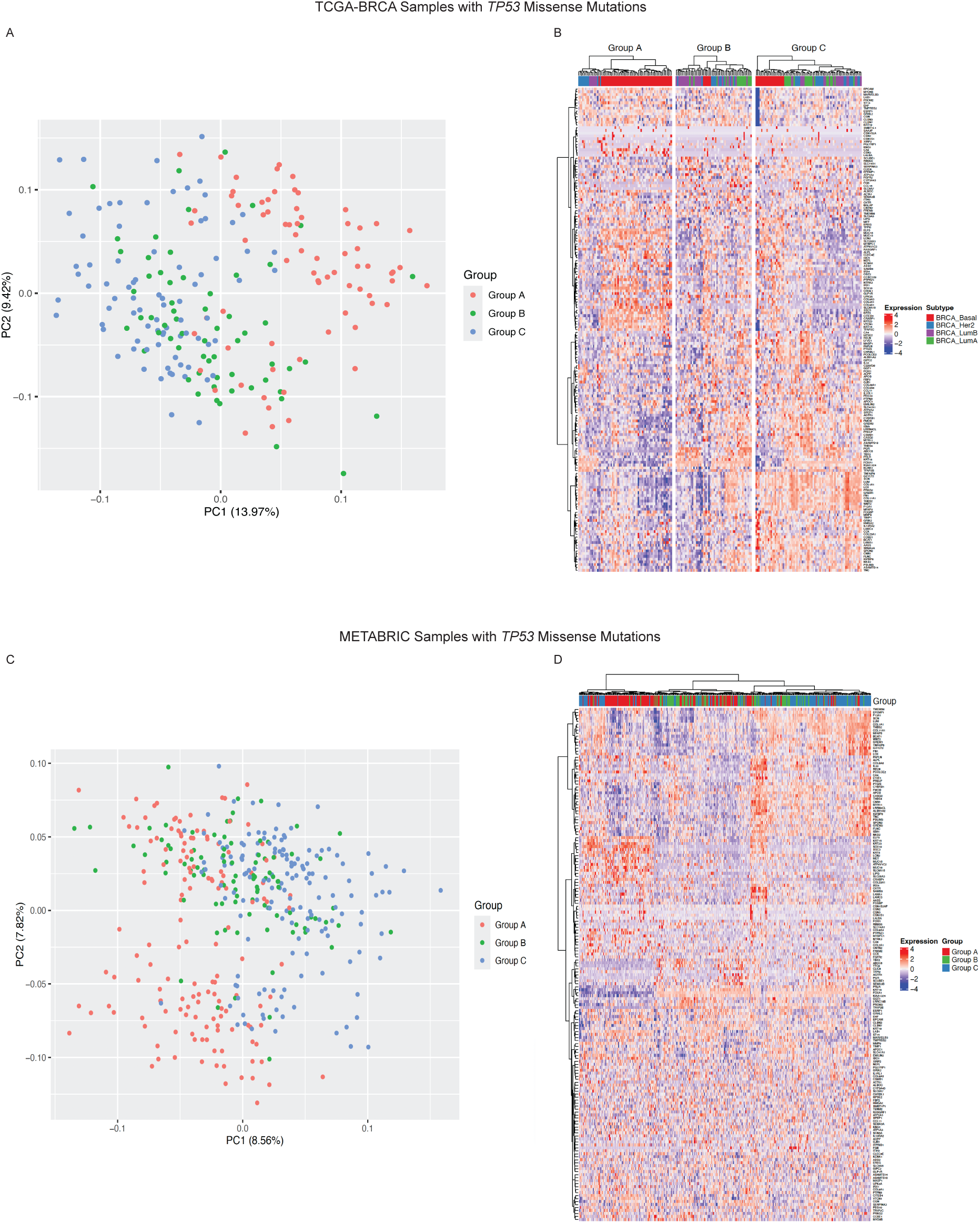
Expression of *MaP^R245W/+^* Intrinsic Signatures in Human Breast Tumors. (A) Principal components analysis depicting the three *MaP^R245W/+^* transcriptional subgroups in human breast tumors with a *TP53* missense mutation catalogued in the TCGA-BRCA dataset (n=214). (B) Heatmap showing supervised hierarchical clustering of TCGA-BRCA tumors with *TP53* missense mutations grouped according to the *MaP^R245W/+^* intrinsic gene signatures. (C) Principal components analysis depicting the three *MaP^R245W/+^* transcriptional subgroups in human breast tumors with a *TP53* missense mutation catalogued in the METABRIC dataset (n=394). (D) Heatmap showing supervised hierarchical clustering of METABRIC tumors with *TP53* missense mutations grouped according to the *MaP^R245W/+^* intrinsic gene signatures.

## Discussion

In this study we have demonstrated that a single initiating p53 mutation gives rise to three molecularly distinct metastatic breast cancer tumor types: (1) Stem Cell Like (2) Well-Differentiated Metabolically Active and (3) Immunosuppressive. Stem cell like tumors are characterized by activation of pathways involving proliferation and ribosomal biogenesis, amplification of *Met*, deletion of *Nf1* and enrichment of resting dendritic cells to promote an immune tolerant environment. Well-differentiated metabolically active tumors enriched for lipid and amino acid metabolic processes, estrogen signaling, recurrent mutations in the *Pten*, *Pik3ca*, and *Pik3r1* to activate Pi3k signaling, and enrichment of CD8^+^ T cells. Immunosuppressive tumors are uniquely distinct from the stem cell like and well-differentiated metabolically active tumors, as they do not activate any hallmark cancer pathways, but rather enrich biological processes reflective of immune suppression. Additionally, these tumors have the highest mutation burden and harbor mutations in *Traf7*, delete *Cdkn2a*, and enrich for M2 macrophages.

Our meta-analyses of human breast tumors in the TCGA-BRCA and METABRIC datasets, demonstrated that tumors harboring *TP53* missense mutations express the intrinsic gene signatures identified in our murine model of mBC. PCA and hierarchical clustering of the human data was not as robust as our murine transcriptomic analyses, however, this is likely because the timing of the *p53* mutation is unknown in these tumors, opposed to it being the *initiating* event in our system. Furthermore, the human breast tumor cohort consists of a mixture *TP53* mutations, in contrast to the homogeneity of our murine cohort. Nonetheless, our human analysis underscored the fact our intrinsic mBC groupings are not restricted to the *TP53^R248W^*mutation nor to a particular molecular subtype of breast cancer. It is well established that *TP53* mutations are most prevalent in TNBC, however, our results corroborate other studies that suggest breast tumors with a *p53* mutation, regardless of their molecular subtype, are likely more aggressive. As a result, further investigation of our intrinsic mBC subtypes is warranted to unveil potential therapeutic opportunities not only in TNBC, but also mBCs of other molecular subtypes harboring a p53 mutation.

Our intrinsic mBC subtypes have group specific molecular signatures that may be vulnerabilities for targeted therapy. Our stem cell-like tumors amplify the *Met* oncogene, which has been demonstrated to mediate metastasis in various cancers (*117*). Tumors with a similar molecular and genomic profile may respond to the MET receptor tyrosine kinase inhibitor, Capmatinib, an FDA approved drug used to treat non-small cell lung cancers with *MET* amplification or MET exon 14 skipping (*118*). Stem cell-like tumors also amplify *Yap1* and *Birc*3. Tumors in this group may also respond to the FDA-approved Yap1-TEAD inhibitor, Verteporfin, which has been shown to have anti-tumor effects in metastatic breast cancers (*119*). These tumors may also be responsive to one or more Smac mimetics or inhibitor of apoptosis (IAP) antagonists, which are currently investigated in clinical trials for their ability to inhibit Birc3 in squamous cell head & neck carcinoma (*120*).

Our well-differentiated metabolically active tumors exclusively activate the PAM signaling pathway via mutations in *Pten*, *Pik3ca*, and *Pik3r1*. Given this genomic profile, these tumors may be responsive to the PI3K inhibitor, Alpelisib, which is currently FDA-approved for hormone receptor negative/HER2+ advanced or metastatic breast cancers with a *PIK3CA* mutation (*121*). Recent clinical trials results demonstrate patients HR+ mBC and mutation of PIK3CA have a 30% overall response rate to Apelisib, while TNBC patients with *PIK3CA* mutations that did not clinically benefit from it (*122*) Another potential treatment option for these tumors, capivasertib, which is an AKT inhibitor that has been approved as a combination therapy in metastatic hormone receptor positive, HER2-negative breast cancers with *PIK3CA* mutation and/or *PTEN* loss (*123*). Early clinical trials testing the efficacy of capivasertib as a treatment for TNBC with *PIK3CA* mutation and/or *PTEN* loss were halted as predefined overall survival goals were not met. While it seems that HR+ breast tumors with *PIK3CA* and/or PTEN alterations respond positively to PI3K pathway inhibitors such as Apelisib and Capivasertib, the *TP53* status is unknown in these tumors. Our murine Group B well-differentiated metabolically active tumors consist of both TNBC and HR+ (luminal B) tumors and thus share a common transcriptional program associated with resistance to these drugs. Therefore, tumors within this group may benefit from therapies targeting transcriptional regulators. Our eQTL analysis demonstrated the transcription factor, Tbx1 (amplified in 46% of tumors) regulates genes involved in ECM and connective tissue, suggesting it may have a role in metastatic progression. A single *in vitro* study has shown TBX1 regulates proliferation and invasion in breast cancer; but it has not been studied *in vivo*, specifically in the context of a p53 mutation. Thus, further studies are warranted to investigate Tbx1 as a therapeutic target in Group B well-differentiated metabolically active breast tumors.

Immunosuppressive tumors in our model have a high TMB and have recurrent mutations that are highly predictable of response to immunotherapy. Tumors in this group mutate the COSMIC oncogene *Traf7.* Meningiomas are similarly immunosuppressive, mutate *Traf7* and have a high TMB. Recent case studies and phase 2 clinical trials demonstrate efficacy of the immunotherapy, pembrolizumab in patients with these aggressive tumors (*124, 125*). Additionally, Group C immunosuppressive tumors recurrently mutate *Ttn*, *Muc4*, and *Ccdc168* which are also associated with response to immunotherapy. This tumor group also mutates *Il4ra*, which mediates immunosuppression. Dupixent is a monoclonal antibody against IL4R and is used to suppress type 2 immune reactions to treat various inflammatory conditions (*126*). Phase I clinical trials are currently underway to assess its utility as a combination therapy to improve response to immunotherapy and/or chemotherapy in lung cancer and TNBC. Thus, breast tumors with a *TP53* missense mutation and expression of our immunosuppressive signature may respond to pembrolizumab and Dupixent combination therapy. Collectively, the human mBCs harboring *TP53* missense mutation and expression of one of our intrinsic signatures may benefit from one or more existing FDA-approved drugs for targeted therapy.

The clinical utility of germline *TP53* mutations in breast cancer is well-established and is important for guiding therapy. However, the clinical utility of somatic missense mutations in *TP53* is not established in breast cancer. This study shows a single *Trp53* mutation induces three intrinsic groups of metastatic breast cancer. The transcriptomic signatures defining each tumor group exist in human breast tumors with a *TP53* missense mutation. Follow up machine learning analyses are necessary to determine the minimal number of genes necessary to generate a gene signature that accurately distinguishes these intrinsic groups in human breast cancer patients. Establishment of such a gene signature will be useful for guiding targeted therapy in mBC patients with a *TP53* missense mutation.

## Materials and Methods

### Mice and tumor harvesting

Female *MaP^R245^* mice in an F1 hybrid 50% BALB/c and 50% C57 BL6/J background, were generated via mammary gland injection of Ad-Cre, at the age of 10-12 weeks, as previously described (*22–24*). All mice were genotyped by polymerase chain reaction (PCR) as previously described (*22–24*). The mouse cohort was monitored daily for tumor development. Once moribund, mice were euthanized according to IACUC guidelines and tissues were collected in 10% v/v formalin and paraffin embedded. A portion of mammary tumors and matched metastases were flash frozen on dry-ice and stored for downstream analyses. All mouse experiments were performed in compliance with the guidelines of the Association for Assessment and Accreditation of Laboratory Animal Care International and the US Public Health Service Policy on Human Care and Use of Laboratory Animals.

### Molecular subtyping

Molecular subtyping of mammary tumors to define *Esr1* (estrogen receptor), *Pgr* (progesterone receptor) and *Erbb2* (HER2) expression was performed via qRT-PCR analysis as previously described (*22–24*).

### Histology

Mammary tissues harvested from mice were fixed in 10% neutral buffered formalin, followed by paraffin embedding. 4 mm tissue sections were stained with hematoxylin and eosin (H&E) by the MD Anderson Cancer Center Department of Veterinary Medicine Surgery and Histology Laboratory. Tissue sections were analyzed by two pathologists. H&E bright field images were taken at 20X using the Nikon Eclipse Ni microscope.

### RNA extraction

Mouse mammary tumors were homogenized using Trizol and Qiagen RNeasy Kits to isolate total RNA using a modified extraction protocol as previously described (*23, 24*). Trizol was added to homogenized tissue and incubated at room temperature for 5 minutes. A 1:5 volume of chloroform was added to the tissue/Trizol mixture and vortexed briefly. Samples were incubated at room temperature for 3 minutes, followed by centrifugation at 12000g at 4°C for 30 minutes. The aqueous phase was transferred to a new collection tube before mixing with 1.5 volume of 100% ethanol and loading onto a RNeasy spin column (Qiagen, CA), Columns were centrifuged at >8000g for 15 seconds. Flow-through for each sample was discarded. Each column was washed with buffer RW1, treated with DNase I, and then washed with buffers RW1 and RPE, respectively. Residual ethanol was dried with a final spin and RNA was eluted in 60 ml of nuclease free water.

### RNA sequencing

Barcoded, Illumina compatible, stranded total RNA libraries were prepared using Illumina Stranded Total RNA Prep Ligation with Ribo-Zero Plus Kit (Illumina). Briefly, 100ng of DNase I treated total RNA was depleted of cytoplasmic and mitochondrial ribosomal RNA (rRNA) using Ribo-Zero Plus. The depleted RNA was then fragmented and reverse transcribed using random hexamers. Deoxyuridine triphosphate (dUTP) was incorporated during second strand synthesis to provide strand specificity.

The ends of the resulting blunt-ended double stranded cDNA fragments were 3’-A tailed, and pre-index anchors were ligated. The products were then purified, and Illumina specific dual indexes were added using 13 cycles of PCR, to create the final cDNA library. The libraries were quantified using the Qubit dsDNA HS Assay Kit and assessed for size distribution using the Agilent TapeStation (Agilent Technologies), then pooled. The pool was quantified by qPCR using the KAPA Library Quantification Kit (KAPA Biosystems). The following samples were sequenced in 75nt PE format on the NextSeq 500 High: YZ-1, YZ-4, YZ-5, YZ-14, YZ-17, YZ-20, YZ-21, and YZ-24. All remaining samples in the cohort were sequenced in three lanes of the NovaSeq6000 S4-200 flow cell using the 100nt PE format.

### Bulk RNA-seq analysis

RNA-seq FASTQ files were processed through FastQC, a quality control tool to evaluate the quality of sequencing reads at both the base and read levels. Samples that passed QC were subsequently analyzed. STAR alignment (*127*) was performed with default parameters to generate RNA-seq BAM files with GRCm39. Aligned reads were summarized at the gene level using STAR (Version 2.7.10b). Gene-level annotation was performed using the GENCODE vM34 annotation, which was downloaded from the GENCODE project (*128*). The raw count data was processed by Deseq2 (*129*) software to identify differentially expressed genes (DEGs) between sample groups. The final p-value was adjusted using the Benjamini and Hochberg method (Benjamini et al. 1995). A threshold gene expression log2 fold change of >=1.0 or <=-1.0 and an FDR q-value of <=0.05 was applied to select the most significant DEGs. Differential expression analysis was further evaluated utilizing various pathway enrichment tools including GSEA (*35*).

### Whole exome sequencing library preparation

DNA was isolated using the Qiagen DNeasy Blood & Tissue kit according to the manufacturer’s instructions. Illumina compatible, dual indexed libraries were prepared from 50ng of enzymatically fragmented, RNase treated genomic DNA using the Twist Library Preparation Kit and the Twist Universal Adapter System, per the manufacturer’s protocol. The indexed libraries were prepared for capture with 6 cycles of PCR amplification using Twist UDI primers, then assessed for fragment size distribution on the 4200 TapeStation High Sensitivity D1000 ScreenTape (Agilent Technologies) and quantified using the Qubit dsDNA HS Assay Kit (ThermoFisher). Libraries’ concentrations were normalized, and equal quantities of the unique dual indexed libraries were combined, 7-8 libraries per pool.

Exon target capture was performed using the Twist Mouse Core Exome kit. Following capture, the exon enriched library pools were amplified using 6 cycles of PCR then assessed for size distribution using the 4200 TapeStation High Sensitivity D1000 ScreenTape (Agilent Technologies) and quantified using the Qubit dsDNA HS Assay Kit respectively. The exon enriched library pools were quantified by qPCR using the KAPA Library Quantification Kit (KAPABiosystems) and then sequenced, in one lane of the NovaSeq6000 S4-300xp flow cell using the 150nt PE format.

### Somatic mutation detection from whole exome sequencing (WES)

FastQC (version 0.11.5) (*130*) was used to check the quality of each FASTQ file. Sequencing reads were aligned to the mouse reference genome using BWA with BWA MEM mode, and somatic mutation calls were carried out based on Genome Analysis Toolkit (GATK) (*131*) best practices. The GATK tools MarkDuplicates, BaseRecalibrator, and ApplyBQSR were used to identify duplicate reads and to detect and correct for patterns of systematic errors in the base quality scores. Samtools (version 1.9) was used to subsample and merge normal samples to generate a pooled normal sample. Somatic mutations were called using GATK MuTect2 v4.1.0.0 for tumors with the pooled normal sample. False positive variants were filtered out using the default parameters of the GATK tools FilterMutectCalls, CollectSequencingArtifactMetrics, and FilterByOrientationBias. The Ensembl Variant Effect Predictor (VEP) was used to functionally annotate somatic mutations including single nucleotide variants and short insertions/deletions. An additional filtering step was applied to remove mutations with variant allele read depth less than 5.

### Copy number alteration detection from WES

Copy number variation was inferred using the latest version of Control-FREEC (*132*). Control-FREEC utilizes input aligned reads (bam files) to construct copy number and B-allele frequency profiles with tumor purity estimation and correction. The profiles are then normalized, segmented and analyzed to assign copy number events to each genomic region. Using a matched normal sample, Control-FREEC discriminates somatic from germline events.

Copy number events were divided into a minimum of two classes based on their observed frequency: focal copy number events much smaller than a chromosome arm, and broad copy number events that span a chromosome arm or entire chromosome. Broad and focal copy number changes occur at markedly different rates and may have unique biological consequences and thus were separately analyzed. A length threshold of 50% of a chromosome arm was used to distinguish between broad and focal events. To remove false positive segments resulting from hyper-segmentation, segments were filtered using an amplitude threshold at a copy-difference of 0.35 (log scale). The frequency of broad copy number changes was calculated. The GISTIC algorithm was used to analyze focal copy number changes and define the key peaks of amplifications and deletions.

### LOH inference from WES data

Control-FREEC was used to to identify Loss of Heterozygosity (LOH) in whole-exome sequencing (WES) data by concurrently analyzing copy number and B-allele frequency (BAF) from matched tumor and normal sample pairs. Control-FREEC integrates the segmented copy number and BAF profiles. Each segment is evaluated using a maximum-likelihood model to predict the most probable genotype. LOH is defined if a region has a significant deviation of the BAF from the expected 0.5 value in the tumor sample, coupled with a corresponding change in the copy number profile. This dual approach aides the software in distinguishing between different types of genomic alterations and confidently call regions where one of the parental alleles has been lost in the tumor cells. The interplay of copy number and B-Allele frequency in LOH detection allows for the differentiation between various LOH scenarios:

- **Deletion-associated LOH:** This is the most straightforward case. A decrease in the copy number ratio (e.g., a value of 1 in a diploid region) will be accompanied by a BAF that is either 0 or 1, indicating the complete loss of one allele.
- **Copy-Neutral LOH (cnLOH):** This is a more subtle event where the copy number remains normal (e.g., diploid), but one parental chromosome region is replaced by a copy of the other. In this scenario, the copy number ratio would be close to 2, but the BAF at heterozygous sites would shift from 0.5 to either 0 or 1. Without BAF analysis, these events would be missed by tools that only consider read depth.
- **Amplification-associated LOH:** In some cases, one allele is amplified while the other is lost. This would be reflected by an increased copy number and a BAF value deviating from 0.5 towards 0 or 1.

### LOH analysis via sanger sequencing

LOH status was investigated by polymerase chain reaction (PCR) analysis using primers near the exon 7 mutation site of *MaP^R245W/+^* mice, followed by sequencing. The primers used to measure LOH in *MaP^R245W/+^* mice have been previously described (*22, 133*): forward: 5′-CGGTTCCCTCCCATGCTA-3 and reverse: 5′-AGCGTTGGGCATGTGGTA-3’. The criteria for categorizing the status of WT *Trp53* alleles are described in the results and in Fig. S6.

### Expression quantitative trait locus analysis of copy number alterations on mRNA

Expression quantitative trait loci (eQTL) analysis was performed to identify *cis* and *trans* effects of copy number alterations on differential gene expression. Specifically, differential gene expression analysis was performed on tumors harboring significant amplifications and deletions compared to those without them. *Cis*-regulatory events were defined as differentially expressed genes (DEGs) located on the same chromosome of a given CNA. T*rans*-regulatory events were defined as DEGs that were not located on the same chromosome as a CNA. GSEA analysis was performed to determined biological and molecular processes associated with cis- and trans-regulated DEGs.

### CIBERSORT

CIBERSORT (Cell-type Identification By Estimating Relative Subsets Of RNA Transcripts) was employed to infer relative proportions of immune cells based on a gene expression-based deconvolution algorithm and the LM22 gene signature using bulk transcriptomic FPKM data (*107*). Correlation matrix plots were produced using the R packages ‘corrplot’ and ‘chart.correlation’.

### Cross-Species Transcriptomic Similarity Analysis

To investigate whether the intrinsic transcriptomic signatures identified in murine *MaP^R245W/+^* breast tumors are conserved in human breast cancer, we performed a comparative analysis using RNA-seq data from The Cancer Genome Atlas Breast Invasive Carcinoma (TCGA-BRCA) cohort (*15*). Only tumors harboring *TP53* missense mutations were included in the analysis (n=214). A curated gene set consisting of 183 murine genes associated with three transcriptionally defined subgroups: Group A (stem cell-like; predominantly triple-negative breast cancer [TNBC]), Group B (well differentiated metabolically active; luminal A and TNBC), and Group C (immunosuppressive-like; TNBC). After converting to human orthologs our analysis included 175 genes. To assign each human tumor to one of the three murine transcriptomic groups, we implemented a similarity scoring approach. For each gene in each group, we compared the direction of expression change in the murine dataset (upregulated or downregulated) with the gene’s expression in the TCGA sample. If the direction matched, a score of 1 was assigned; if not, a score of 0 was given. This process was repeated for all genes in each group across all tumors. A final similarity score for each group was calculated by summing the total number of matches and dividing by the number of genes in the corresponding gene set. Each human tumor was then assigned to the group for which it had the highest normalized similarity score.

Heatmaps were generated using the ComplexHeatmap R package (version 2.24.1) (*134*). Gene expression values were z-scored by gene, and rows were ordered by murine group classification (Groups A, B, and C). Columns represented human tumors, annotated by PAM50 breast cancer subtype. For hierarchical clustering of both rows and columns, **Manhattan distance** was used to compute pairwise dissimilarities, and clustering was performed using the **Ward.D linkage method**.

This analysis was repeated using RNA-seq data from 394 human breast tumors harboring a *TP53* missense mutation in the METABRIC cohort (*116*). METABRIC tumors were not annotated by PAM50 breast cancer subtypes, as subtype data is not available for tumors in this cohort.

### Statistical analysis

Kaplan-Meier survival analyses for each mouse cohort were performed using Prism 9 software (GraphPad Software, version 9, RRID:SCR_002798, CA). Fisher’s Exact tests comparing primary tumor incidence were performed using Prism 9 software. Statistical analysis of changes in gene expression was performed using the Bioconductor packages *DESeq2* in the R statistical computing environment (Version 4.2.0).

## Supporting information

Supplementary Figures S1-S8; Supplementary Tabls S1-S11

## Author contributions

Conceptualization: JMM, GL

Data curation: JMM, XS, ZVR

Formal analysis: JMM, RLM, VC, XS, ZVR, GPC, LH, AKE

Investigation: JMM, RLM, GPC

Methodology: GL, JMM

Project Administration: GL, JMM

Resources: GL, XS, AK

Software: XS, ZVR

Visualization: JMM, XS, AK, ZVR

Writing—original draft: JMM and GL

Writing-review & editing: JMM, GL, AK

## Data and materials availability

RNA-seq and whole exome seq data reported in this article have been deposited in the Gene Expression Omnibus database, GSE301299 and GSE301300.

