## Supplementary Figures S1-S8; Supplementary Tabls S1-S11 for "Multi-omics analysis identifies intrinsic *Trp53* driven metastatic breast cancer subtypes"

### Multi-omics analysis identifies intrinsic Trp53 driven metastatic breast cancer subtypes—14pt. bold No Paragraph Breaks

Joy Marie McDaniel *et al*.

**This PDF file includes:**

Supplementary Text
Figs. S1 to S8
Tables S1 to S11
Data S1 to S9

#### Supplementary Text

Supplementary Figures and Tables

Supplementary figures and tables include IVIS experiments to characterize the specificity of recombination in the model; histological and molecular characterization of the model; Sanger sequencing results for selected tumors to measure loss of heterozygosity; B-allele frequency plots for loss of heterozygosity inference for individual tumors, and summary of copy number variation across all tumors.

Supplementary Data

Supplementary data includes nonsynonymous mutation data for each tumor subgroup; group specific copy number alterations; and group specific copy number-mRNA eQTL analysis.

Fig. S1.

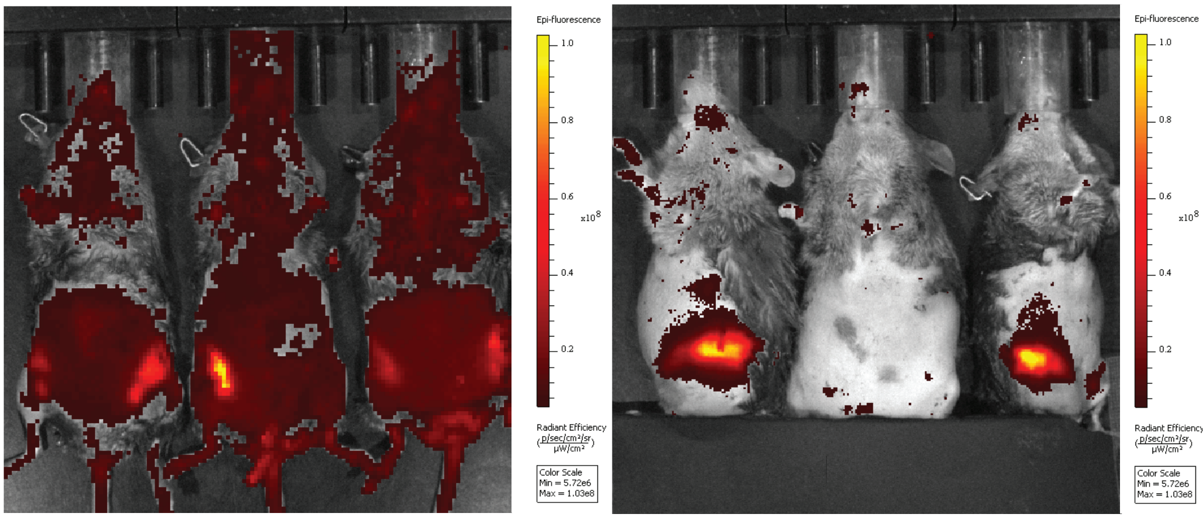


JM956 3 MPI | JM729 7 MPI | JM595 10 MPI | JM409 12 MPI | JM596 10 MPI Negative Control | JM1039 1 MPI

**IVIS Imaging Traces Recombination in *MaP*$^{R245W/+}$ Mice** *MaP*$^{R245W/+}$ mice were subjected to IVIS imaging in the University of Texas MD Anderson Cancer Center Small Animal Imaging Facility (SAIF). Imaging was conducted in the red fluorescence channel to identify TdTomato positive signal in the mammary gland. The color corresponds to fluorescent signal level, and the scale is provided to the right of each image. Below each image from left to right is the mouse ID for each animal, as well as the time elapsed since the mouse was injected with Adenovirus-packaged Cre recombinase. Each timepoint is in months post-injection (MPI). JM596 served as the negative control, as this mouse did not possess the *TdTomato* reporter allele, and therefore shows no TdTomato fluorescence. Fluorescence signals were visible only in the mammary tissue.

**Fig. S2.**

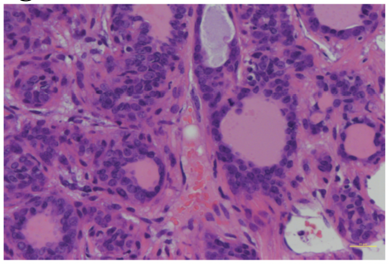

Carcinoma with Adenosis Pattern (JM1384)

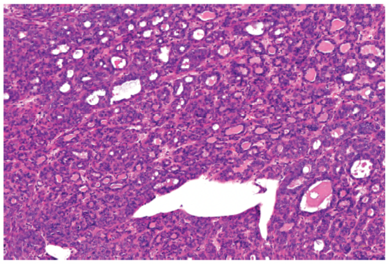

Carcinoma with Papillary Pattern (JM964)

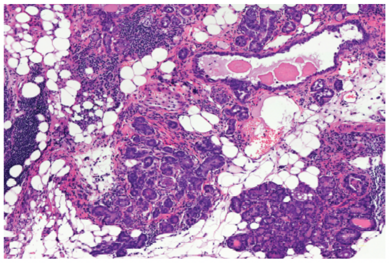

Adenocarcinoma (JM1566-T2)

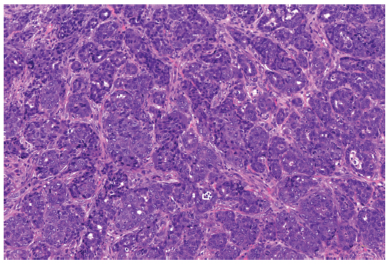

Poorly Differentiated Carcinoma w/Basaloid Features (RM247)

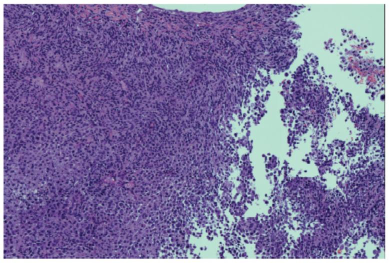

Carcinoma (RM221)

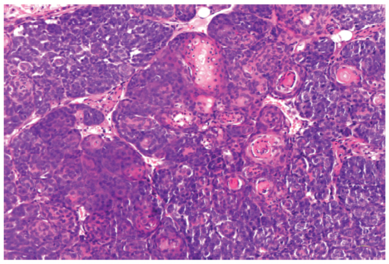

Carcinoma with Squamoid Features (JM1039)

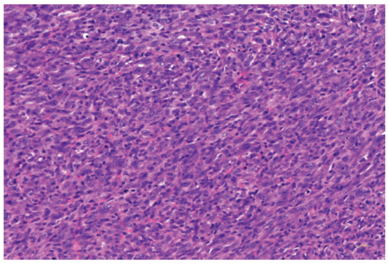

Metaplastic Carcinoma with Spindle Cells (JM1419)

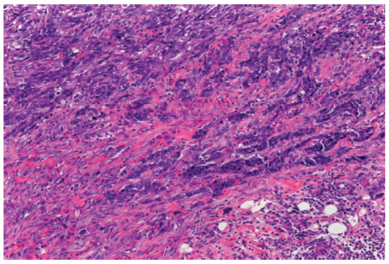

Sarcomatoid Carcinoma with Sarcoma-like & Metaplastic Features (JM884)

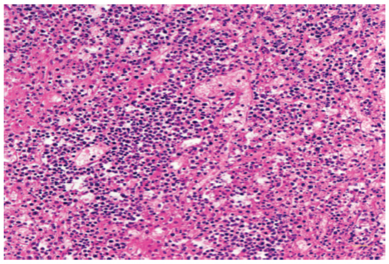

Small Blue Cell Tumor (JM590-T2)

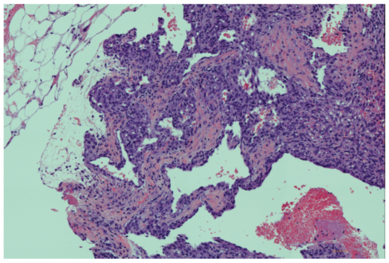

Unclassified (JM731)

**Histology Observed in *MaP*$^{R245W/+}$ Mammary Tumors** H&E images representative of the histological classifications observed in *MaP*$^{R245W/+}$ mammary tumors (20x)

Fig. S3.

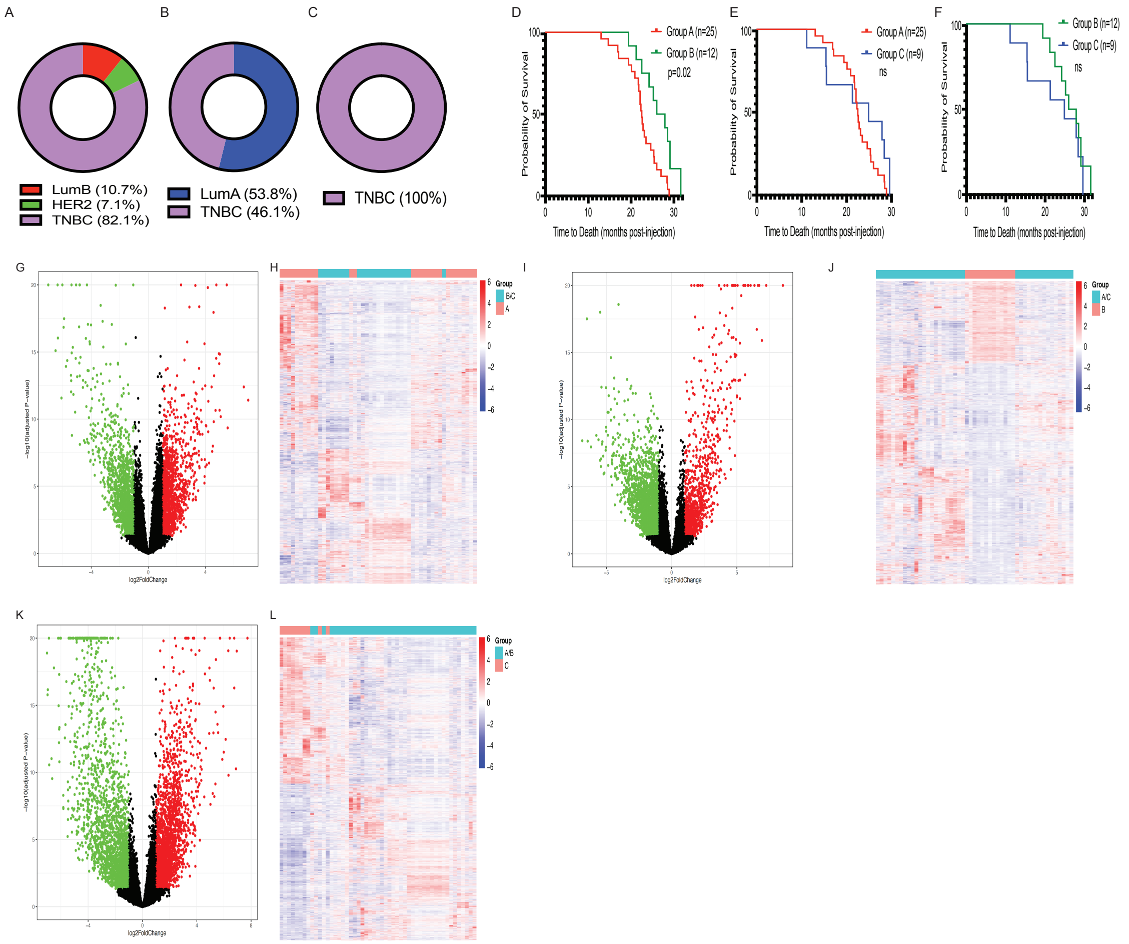


**Additional Molecular Characteristics of *MaP*$^{R245W/+}$ Mammary Tumors** (A-C) Pie charts depicting breast cancer molecular subtypes present in Groups A, B and C mammary tumors. (D-F) Overall survival of Groups A, B, and C *MaP*$^{R245W/+}$ mice. (Intergroup survival comparison: Groups A vs B, p< 0.02; A vs C, ns; B vs C, ns) (G) Volcano plots depicting 1,439 genes upregulated and 1,650 genes downregulated in Group A mammary tumors compared to Groups B/C mammary tumors; (false discovery rate (FDR) of 5% and log2 fold change of |1|). (H) Heatmap showing unsupervised hierarchical clustering of Group A mammary tumors compared to Groups B and C mammary tumors using the Pearson distance and Ward linkage. (I) Volcano plots depicting 988 genes upregulated and 2,204 genes downregulated in Group B mammary tumors compared to Groups A/C mammary tumors (false discovery rate (FDR) of 5% and log2 fold change of |1|). (J) Heatmap showing unsupervised hierarchical clustering of Group B mammary tumors compared to Groups A and C mammary tumors using the Pearson distance and Ward linkage. (K) Volcano plots depicting 2,111 genes upregulated and 2,227 genes downregulated in Group C mammary tumors compared to Groups A/B mammary tumors (false discovery rate (FDR) of 5% and log2 fold change of |1|). (L) Heatmap showing unsupervised hierarchical clustering of Group C mammary tumors compared to Groups A and B mammary tumors using the Pearson distance and Ward linkage.

Fig. S4.

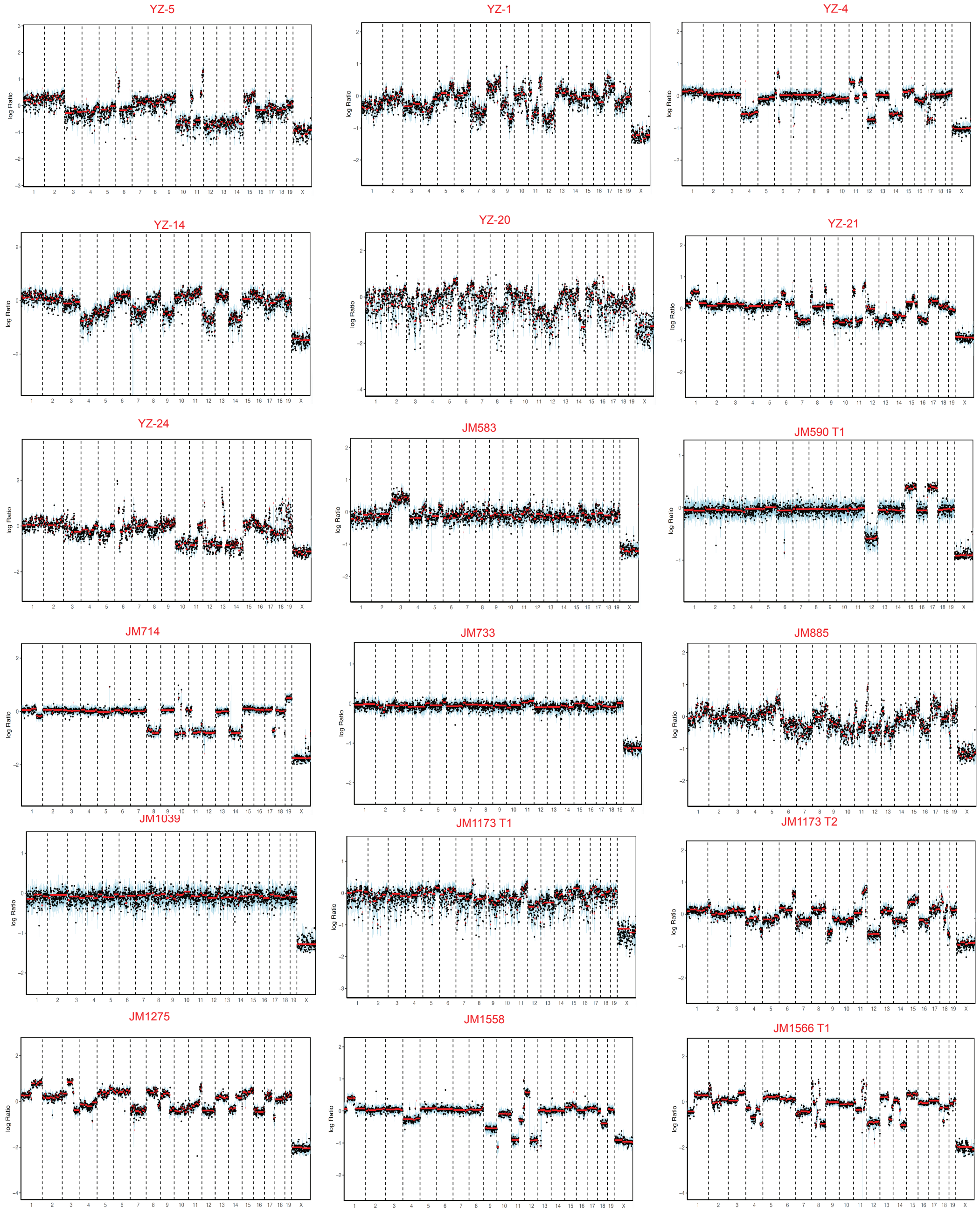


**Inferring LOH from WES Data in Group A Mammary Tumors**

Log ratio plots of B-allele frequency of SNPs in Group A Tumors

Fig. S5.

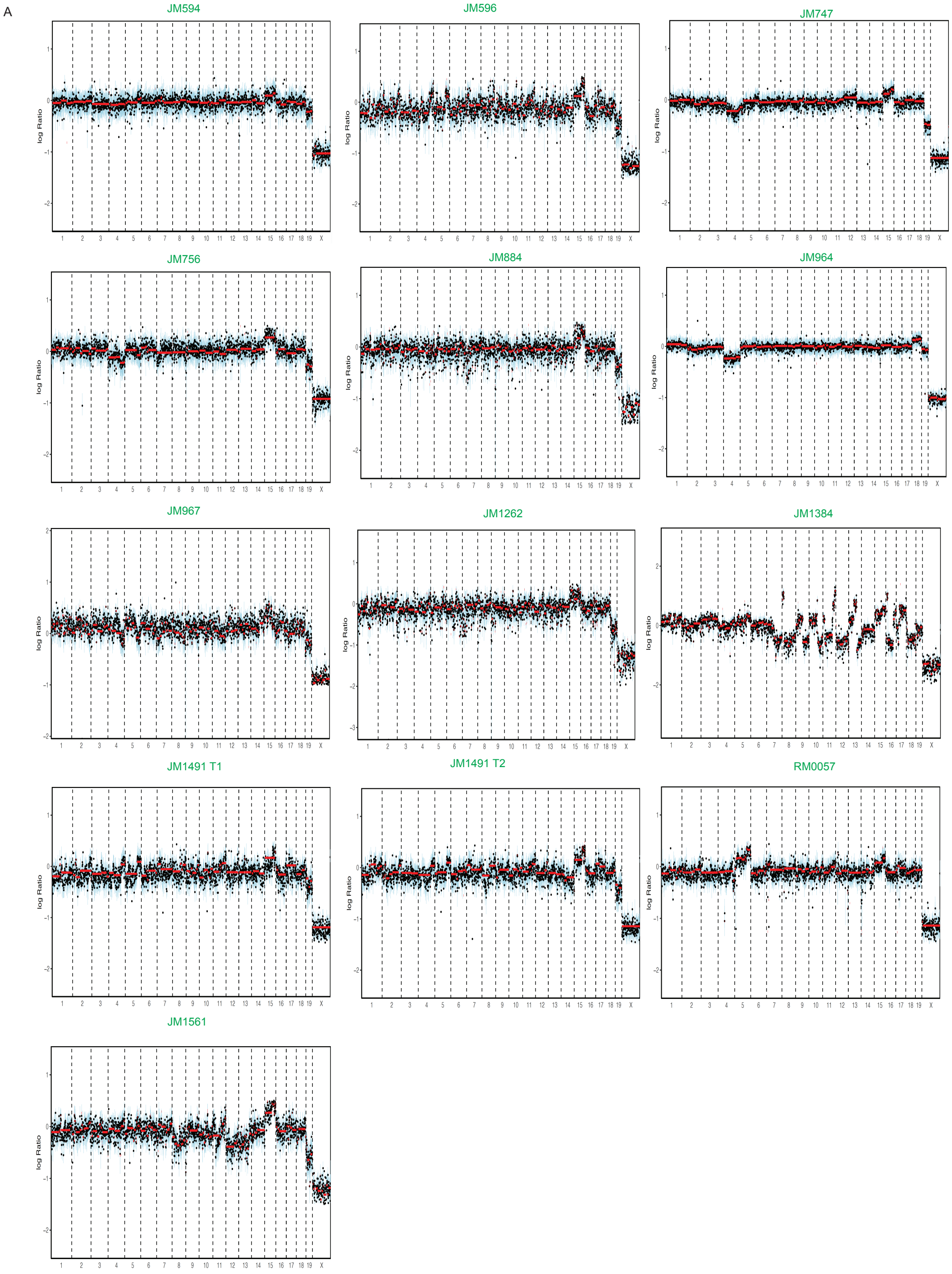


**Inferring LOH from WES Data in Group B Mammary Tumors**

Log ratio plots of B-allele frequency of SNPs in Group B Tumors

Fig. S6.

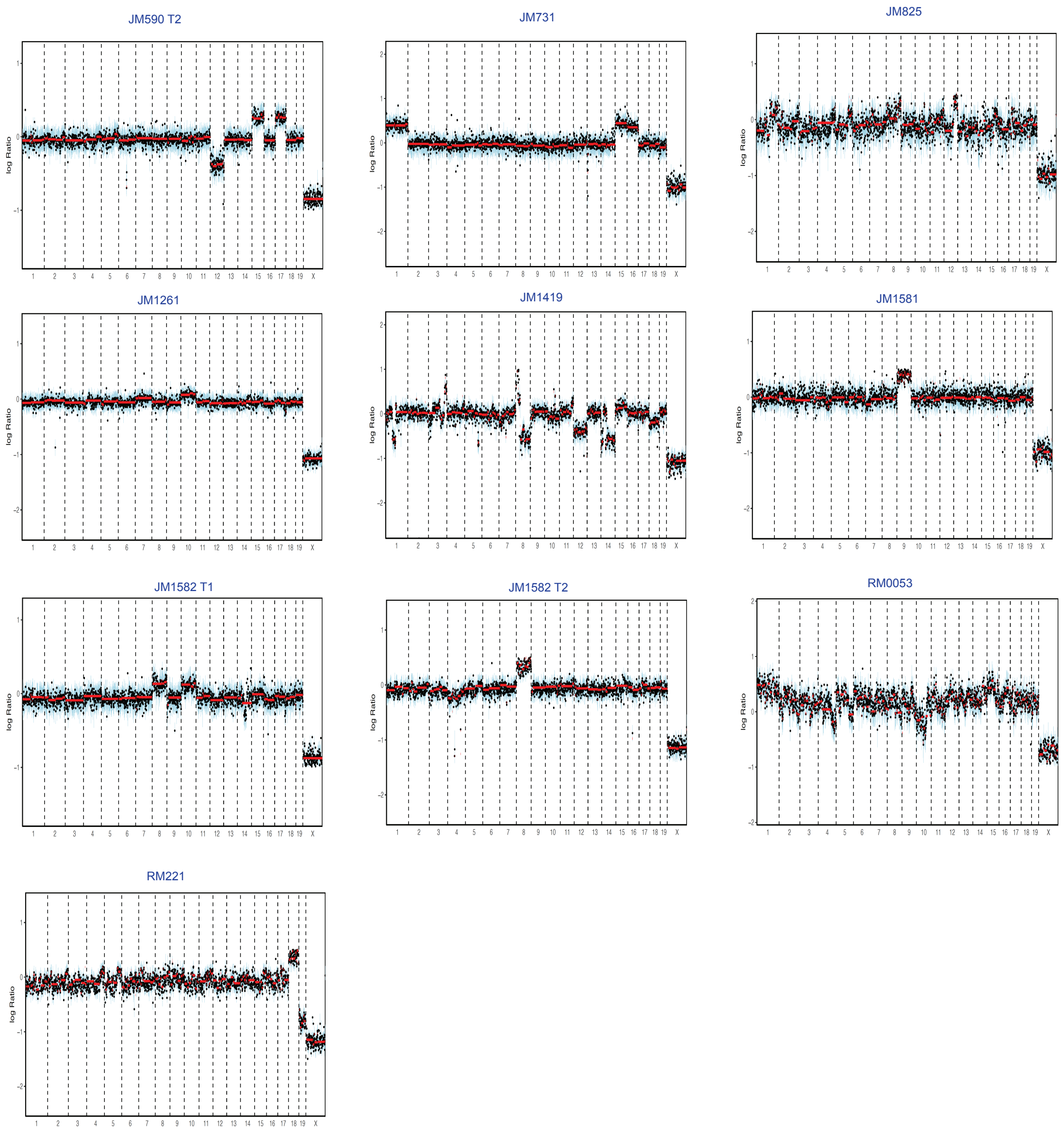


**Inferring LOH from WES Data in Group C Mammary Tumors**

Log ratio plots of B-allele frequency of SNPs in Group C Tumors

Fig. S7.

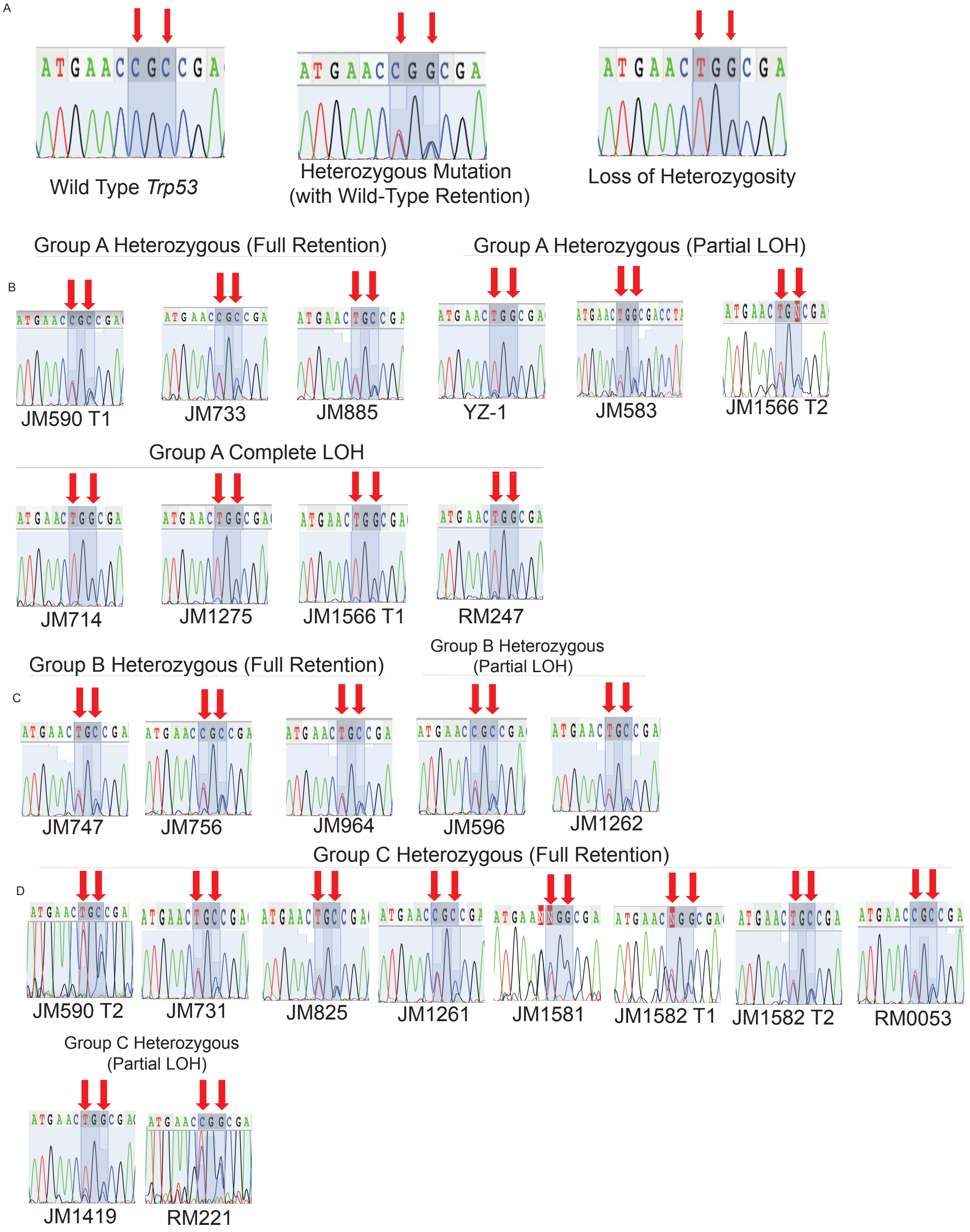


**Sanger Sequencing Characterization of Loss of Heterozygosity at the *Trp53* Locus**
(A)Chromatograms representing wild-type *Trp53*, Heterozygous Mutation of Trp53 with retention of the wild-type allele, and loss of wild type allele (loss heterozygosity) from Sanger sequencing controls (B) Sanger sequencing results for select Group A mammary tumors; (C)

Sanger sequencing results for select Group B mammary tumors (D) Sanger sequencing results for select Group C mammary tumors

Fig. S8.

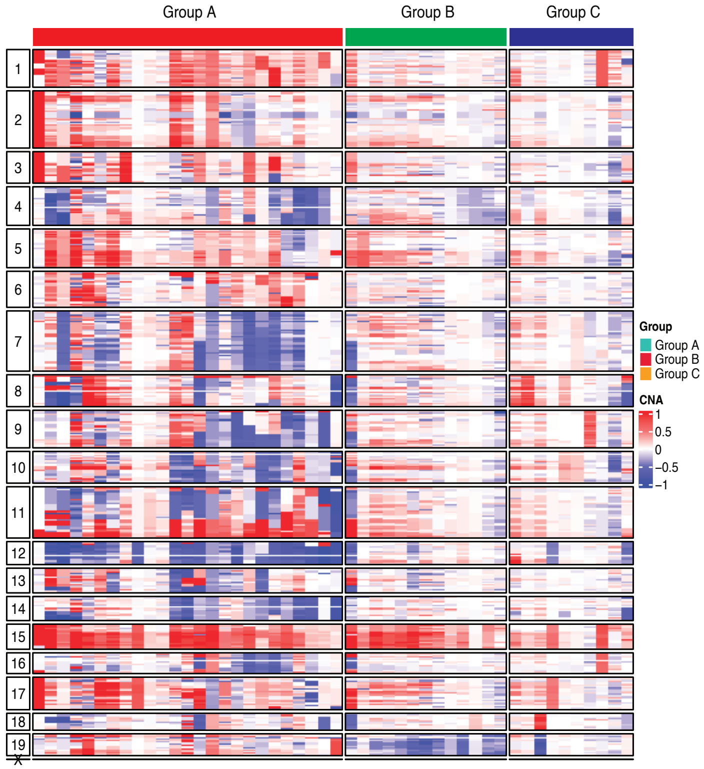


**Copy Number Alterations in *MaP*$^{R245W/+}$ Mammary Tumors**

Heatmap depicting genome-wide copy number alterations inferred from WES of *MaP*$^{R245W/+}$ mammary tumors

Table S1.

| Mouse ID | Histological Classification | Molecular Subtype | Transcriptomic Subgroup | Metastasis | Latency (months post-injection) |
|---|---|---|---|---|---|
| YZ14 | Poorly differentiated carcinoma w/basaloid features | TNBC | Group A | Lung | 14.7 |
| YZ17 | Adenocarcinoma w/angiomatosis | TNBC | Group A | Lung | 21.7 |
| YZ-20 | Adenocarcinoma w/possible matrix (metaplastic) | TNBC | Group A | No | 22.1 |
| YZ-24 | Poorly differentiated carcinoma w/basaloid features | TNBC | Group A | No | 22.1 |
| YZ4 | Poorly differentiated carcinoma w/basaloid features | HER2 | Group A | Lung | 19.4 |
| JM1173-T1 | Adenocarcinoma | TNBC | Group A | Lung | 21.5 |
| JM1173-T2 | Carcinoma (possibly metaplastic) | HER2 | Group A | | |
| JM1381 | Carcinoma w/squamoid features | TNBC | Group A | Liver | 25.6 |
| JM1558 | Adenocarcinoma | TNBC | Group A | Lung | 28.1 |
| JM1566-T1 | Adenocarcinoma | TNBC | Group A | Lung | 22.5 |
| JM1566-T2 | Adenocarcinoma/differentiated carcinoma w/sebaceous features | TNBC | Group A | | |
| JM583 | Poorly differentiated carcinoma w/basaloid features | TNBC | Group A | Lung & Liver | 23.5 |
| RM0008-T1-Cut 1 | Sarcomatoid carcinoma w/sarcoma-like & metaplastic features | TNBC | Group A | Lung | 22.5 |
| RM0008-T1-Cut 2 | Carcinoma w/papillary pattern | TNBC | Group A | | |
| RM0058 | Metaplastic carcinoma, spindle cells | TNBC | Group A | Lung | 22.3 |
| RM247 | Poorly differentiated carcinoma w/basaloid features | TNBC | Group A | Brain, Lung & Skeletal Muscle | 12.8 |
| RM249 | Adenocarcinoma | TNBC | Group A | Lung & Liver | 19.8 |
| RM0087 | Carcinoma w/squamoid features | TNBC | Group A | Lung | 22.5 |
| YZ5 | Sarcomatoid carcinoma w/sarcoma-like & metaplastic features | LumB | Group A | Lung & Liver | 16.9 |
| YZ-1 | Carcinoma w/adenosis pattern | LumB | Group A | No | 17.1 |
| YZ21 | Adenocarcinoma | LumB | Group A | Lung | 23.1 |
| JM1275 | Metaplastic carcinoma, spindle cells | TNBC | Group A | No | 20.6 |
| JM590-T1 | Small blue cell tumor | TNBC | Group A | Lung & Liver | 22.5 |
| JM733 | Adenocarcinoma/differentiated carcinoma (glands) w/lymphoid tissue | TNBC | Group A | Lung & Ovary | 16.5 |
| JM885 | Sarcomatoid carcinoma w/sarcoma-like & metaplastic features | TNBC | Group A | No | 22 |
| JM1039 | Carcinoma w/squamoid features | TNBC | Group A | Lung & Liver | 24.9 |
| JM1193 | Unclassified | TNBC | Group A | Lung | 22.5 |
| JM714 | Adenocarcinoma | TNBC | Group A | No | 25 |
| JM1491-T1 | Carcinoma w/adenosis pattern | LumA | Group B | Left Eye Growth (Brain) | 22.5 |
| JM1491-T2 | Carcinoma w/adenosis pattern | TNBC | Group B | | |
| JM1262 | Carcinoma w/adenosis pattern | LumA | Group B | Lung | 20.9 |
| JM1384 | Carcinoma w/adenosis pattern | LumA | Group B | Lung, Liver, Ovary, & Brain | 28.7 |
| JM1561 | Peripheral carcinoma w/papillary pattern | TNBC | Group B | Lung | 22.5 |
| JM594 | Carcinoma w/papillary pattern | LumA | Group B | Lung | 19.5 |
| JM596 | Carcinoma w/adenosis pattern | TNBC | Group B | Lung | 22.5 |
| JM747 | Carcinoma w/adenosis pattern | TNBC | Group B | Lung, Liver & Brain | 22.5 |
| JM756 | Carcinoma w/adenosis pattern | LumA | Group B | Lung | 31.2 |
| JM884 | Carcinoma w/adenosis pattern | TNBC | Group B | Lung & Liver | 27.5 |
| JM964 | Carcinoma w/papillary pattern | LumA | Group B | Lung | 22.5 |
| JM967 | Carcinoma w/adenosis pattern | LumA | Group B | Lung & Liver | 25.8 |
| RM0057 | Carcinoma w/adenosis pattern | TNBC | Group B | Lung | 22.5 |
| JM1419 | Metaplastic carcinoma, spindle cells | TNBC | Group C | Lung | 9.9 |
| JM590-T2 | Small blue cell tumor | TNBC | Group C | Lung & Liver | 22.5 |
| JM1261 | Metaplastic carcinoma, spindle cells | TNBC | Group C | Lung & Liver | 24.6 |
| JM1581 | Metaplastic carcinoma, spindle cells | TNBC | Group C | Lung | 27.6 |
| JM1582-Cut 1 | Metaplastic carcinoma, spindle cells | TNBC | Group C | Lung & Liver | 29.2 |
| JM1582-Cut 2 | Metaplastic carcinoma, spindle cells | TNBC | Group C | | |
| JM825 | Sarcomatoid carcinoma w/sarcoma-like & metaplastic features | TNBC | Group C | Lung, Liver, & Brain | 29.2 |
| RM0053 | Metaplastic carcinoma, spindle cells | TNBC | Group C | Lung & Ovaries | 15.1 |
| RM221 | Carcinoma | TNBC | Group C | Lung & Liver | 24.3 |
| JM731 | Unclassified | TNBC | Group C | Lung | 21 |

**Mice Studied**

#Tumors RNA-Sequenced

▲Tumors Whole Exome Sequenced

Table S2.

| Tumor ID | Group | C:T Ratio (Sanger) | Sanger LOH Status (Sanger) | Allele Copies | Genotype (WES) | LOH Status (WES) | Tumor Mutation Burden (Median) | Transition or Transversion Enriched |
|---|---|---|---|---|---|---|---|---|
| YZ-5 | Group A | N/A | N/A | 2 | AB | HET | 1.3 | Transition |
| YZ-1 | Group A | 24.20% | Partial LOH | 2 | AB | HET | 1.3 | Transition |
| YZ-4 | Group A | N/A | N/A | 2 | AB | HET | 1.3 | Transition |
| YZ-14 | Group A | N/A | N/A | 4 | AABB | HET | 1.3 | Transition |
| YZ-20 | Group A | N/A | N/A | 2 | BB | LOH | 1.3 | Transition |
| YZ-21 | Group A | N/A | N/A | 2 | BB | WT | 1.3 | Transition |
| YZ-24 | Group A | N/A | N/A | 2 | BB | LOH | 1.3 | Transition |
| JM583 | Group A | 71% | Partial LOH | 4 | AABB | HET | 1.3 | Transition |
| JM590 T1 (L45) | Group A | 80% | HET (Full Retention) | 2 | AB | HET | 1.3 | Transition |
| JM714 | Group A | 9.35% | Full LOH | 2 | BB | LOH | 1.3 | Transition |
| JM733 | Group A | 92.60% | HET (Full Retention) | 2 | AB | HET | 1.3 | Transition |
| JM885 | Group A | 87.60% | HET (Full Retention) | 2 | AB | HET | 1.3 | Transition |
| JM1039 | Group A | N/A | N/A | 2 | AB | HET | 1.3 | Transition |
| JM1173 T1 (L45) | Group A | N/A | N/A | 2 | AB | HET | 1.3 | Transition |
| JM1173 T2 (R45) | Group A | N/A | N/A | 2 | AB | HET | 1.3 | Transition |
| JM1275 | Group A | 13.80% | Full LOH | 3 | BBB | LOH | 1.3 | Transition |
| JM1558 | Group A | N/A | N/A | 15 | ABBBBBBBBBBBBBB | HET | 1.3 | Transition |
| JM1566 T1 | Group A | 11.20% | Full LOH | 2 | AB | HET | 1.3 | Transition |
| JM1566 T2 | Group A | 69.20% | Partial LOH | 2 | AB | HET | 1.3 | Transition |
| RM008 BT C2 | Group A | N/A | N/A | 2 | AB | HET | 1.3 | Transition |
| RM008 BT C1 | Group A | N/A | N/A | 2 | AB | HET | 1.3 | Transition |
| RM0058 | Group A | N/A | N/A | 3 | BBB | LOH | 1.3 | Transition |
| RM0087 | Group A | N/A | N/A | 2 | AB | HET | 1.3 | Transition |
| RM247 | Group A | 17.70% | Full LOH | 2 | BB | LOH | 1.3 | Transition |
| RM249 | Group A | N/A | N/A | 3 | BBB | LOH | 1.3 | Transition |
| JM594 | Group B | N/A | N/A | 2 | AB | HET | 1.3 | Transition |
| JM596 | Group B | 77.80% | Partial LOH | 2 | AB | HET | 1.3 | Transition |
| JM747 | Group B | 83.30% | Het (Full Retention) | 2 | AB | HET | 1.3 | Transition |
| JM756 | Group B | 84.70% | Het (Full Retention) | 2 | AB | HET | 1.3 | Transition |
| JM884 | Group B | N/A | N/A | 2 | AB | HET | 1.3 | Transition |
| JM964 | Group B | 86.70% | Het (Full Retention) | 2 | AB | HET | 1.3 | Transition |
| JM967 | Group B | N/A | N/A | 2 | AB | HET | 1.3 | Transition |
| JM1262 | Group B | 76.90% | Partial LOH | 2 | AB | HET | 1.3 | Transition |
| JM1384 | Group B | N/A | N/A | 2 | AB | HET | 1.3 | Transition |
| JM1491 T1 | Group B | N/A | N/A | 2 | AB | HET | 1.3 | Transition |
| JM1491 T2 | Group B | N/A | N/A | 2 | AB | HET | 1.3 | Transition |
| JM1561 | Group B | N/A | N/A | 2 | AB | HET | 1.3 | Transition |
| RM0057 | Group B | N/A | N/A | 2 | AB | HET | 1.3 | Transition |
| JM590 T2 (R45) | Group C | 89.30% | HET (Full Retention) | 2 | AB | HET | 3.3 | Transversion |
| JM731 | Group C | 87.9 | HET (Full Retention) | 2 | AB | HET | 3.3 | Transversion |
| JM825 | Group C | 87.50% | HET (Full Retention) | 2 | AB | HET | 3.3 | Transversion |
| JM1261 | Group C | 100% | HET (Full Retention) | 2 | AB | HET | 3.3 | Transversion |
| JM1419 | Group C | 100% | HET (Full Retention) | 2 | AB | HET | 3.3 | Transversion |
| JM1581 | Group C | 84.10% | HET (Full Retention) | 2 | AB | HET | 3.3 | Transversion |
| JM1582 T1 | Group C | 89.50% | HET(Full Retention) | 2 | AB | HET | 3.3 | Transversion |
| JM1582 T2 | Group C | 86.80% | HET (Full Retention) | 2 | AB | HET | 3.3 | Transversion |
| RM0053 | Group C | 40.70% | Partial LOH | 2 | AB | HET | 3.3 | Transversion |
| RM221 | Group C | 78.70% | Partial LOH | 2 | AB | HET | 3.3 | Transversion |

Summary of LOH Status Across *MaP*$^{R245W/+}$ Mammary Tumors

Red: Group A Stem Cell Like Tumors.

Green: Group B Well-Differentiated Metabolic Active Tumors.

Blue: Group C Immune Suppressed Tumors.

Data S1. (separate file).
Nonsynonymous Mutations in Group A *MaP*$^{R245W/+}$ Tumors

Data S2. (separate file).
Nonsynonymous Mutations in Group B *MaP*$^{R245W/+}$ Tumors

Data S3. (separate file).
Nonsynonymous Mutations in Group C *MaP*$^{R245W/+}$ Tumors

Data S4. (separate file).
Summary of Group A Copy Number Alterations

Data S5. (separate file).
Group A CNA-mRNA eQTL Analysis

Data S6. (separate file).
Summary of Group B Copy Number Alterations

Data S7. (separate file).
Group B CNA-mRNA eQTL Analysis

Data S8. (separate file).
Summary of Group C Copy Number Alterations

Data S9. (separate file).
Group C CNA-mRNA eQTL Analysis
